# Quantifying the contribution of Neanderthal introgression to the heritability of complex traits

**DOI:** 10.1101/2020.06.08.140087

**Authors:** Evonne McArthur, David Rinker, John A. Capra

## Abstract

**Background:** Nearly all Eurasians have ∼2% Neanderthal ancestry due to several events of inbreeding between anatomically modern humans and archaic hominins. Previous studies characterizing the legacy of Neanderthal ancestry in modern Eurasians have identified examples of both adaptive and deleterious effects of admixture. However, we lack a comprehensive understanding of the genome-wide influence of Neanderthal introgression on modern human diseases and traits.

**Results:** We integrate recent maps of Neanderthal ancestry with well-powered association studies for more than 400 diverse traits to estimate heritability enrichment patterns in regions of the human genome tolerant of Neanderthal ancestry and in introgressed Neanderthal variants themselves. First, we find that variants in regions tolerant of Neanderthal ancestry are depleted of heritability for all traits considered, except skin and hair-related traits. Second, the introgressed variants remaining in modern Europeans are depleted of heritability for most traits; however, we discover that they are enriched for heritability of several traits with potential relevance to human adaptation to non-African environments, including hair and skin traits, autoimmunity, chronotype, bone density, lung capacity, and menopause age. To better understand the phenotypic consequences of these enrichments, we adapt recent methods to test for consistent directional effects of introgressed alleles, and we find directionality for several traits. Finally, we use a direction-of-effect-aware approach to highlight novel candidate introgressed variants that influence risk for disease.

**Conclusion:** Our results demonstrate that genomic regions retaining Neanderthal ancestry are not only less functional at the molecular-level, but are also depleted for variation influencing a diverse array of complex traits in modern humans. In spite of this depletion, we identify traits where introgression has an outsized effect. Integrating our results, we propose a framework for using quantification of trait heritability and direction of effect in introgressed regions to understand how Neanderthals were different from modern humans, how selection acted on different traits, and how introgression may have facilitated adaptation to non-African environments.

## INTRODUCTION

Anatomically modern humans (AMH) interbred with archaic hominin groups on multiple occasions and in several locations over the past 50,000 years. As a result, nearly all Eurasians have ∼2% Neanderthal ancestry resulting from interbreeding events that occurred shortly after their ancestors left Africa.^1,2^ Analyses of available genome-wide association studies and large-scale biobank data revealed that alleles of Neanderthal ancestry are associated with diverse traits in modern Eurasians.^1,3–5^ However, due to the limited phenotype data and technical challenges quantifying associations between archaic alleles and traits, previous studies have not comprehensively characterized the genome-wide influence of Neanderthal introgression on modern human diseases and traits.

Archaic admixture may have facilitated the ability of AMH to inhabit diverse environments as they spread around globe.^6^ Several archaic alleles have functions and evolutionary signatures suggestive of positive selection potentially due to beneficial effects in AMH.^6–8^ Many of these alleles influence systems that directly interact with the environment, such as the immune system^9–16^, hair and skin^17–19^, response to oxygen^20^, and metabolism.^7,21,22^

In spite of these potential adaptive benefits of admixture, theory and simulations suggest that introgressed alleles were largely deleterious in AMH.^23,24^ Accordingly, several lines of evidence demonstrate selection against introgressed Neanderthal DNA in most functional regions of human genomes. First, Neanderthal ancestry is depleted in regions of the genome with strong background selection and evolutionary conservation.^17,18,25^ Second, Neanderthal ancestry is depleted in regions of the genome with annotated molecular functions (*e.g*., genes and gene regulatory elements), and this depletion is strongest in annotated brain and testis regulatory regions.^25–27^ Third, remaining alleles of Neanderthal ancestry—*i.e.* introgressed alleles that were maintained by either selection or drift since admixture—are predicted to have different effects on protein and regulatory function than matched sets of alleles that arose on the human lineage.^5,28^ The vast majority of archaic alleles that are strongly associated with disease are risk increasing in the context of modern human populations.^3^

Several non-exclusive scenarios may explain the apparent genetic cost of Neanderthal introgression. Neanderthals had significantly smaller effective population size than AMH populations; thus, less effective selection resulted in the accumulation of weakly deleterious alleles in Neanderthal populations.^29^ After introgression, these variants were subject to more effective selection in larger AMH populations.^23,24^ It is also possible that hybrid incompatibilities and deleterious epistatic interactions between Neanderthal and AMH alleles reduced the fitness of early hybrids.^17,18,26,30,31^

Given the overall negative selection against alleles of Neanderthal ancestry in functional regions coupled with the examples of positive selection on specific introgressed Neanderthal alleles, there is a need to more comprehensively characterize and reconcile the functional effects of introgressed alleles on variation in diverse AMH traits. Previously, the legacy of introgression in AMHs has been largely characterized based on enrichment for molecular annotations (*e.g*., gene function or regulatory element activity)^17,18,25,26^ or existing genome-wide association study (GWAS) hits.^1,3,4^ However, most medically- and evolutionarily-relevant traits are complex, with hundreds or thousands of loci across the genome contributing to them.^32,33^ Thus, studies of individual loci are not sufficient to address the overall influence of Neanderthal admixture on human traits.

Here, we leverage recent maps of Neanderthal ancestry^34^ with powerful new techniques to characterize heritability of common complex traits associated with Neanderthal introgression and identify Neanderthal variant directions of effect.^35,36^ Using well-powered GWASs for more than 400 diverse traits from existing studies and the UK Biobank^37^, we estimate trait heritability over all variation in regions of the human genome tolerant of Neanderthal ancestry and in introgressed Neanderthal alleles themselves. This comprehensive view of the influence of Neanderthal ancestry genome-wide supports selection against Neanderthal ancestry in regions of the genome that influence nearly all complex traits. However, it reveals that introgressed Neanderthal alleles have an outsized effect on several traits with potential relevance for AMH adaptation into non-African environments.

## RESULTS

### Genomic regions tolerant of Neanderthal ancestry are broadly depleted of complex trait heritability

To quantify the relationship between heritability of complex traits and Neanderthal introgression, we first investigated genomic regions where Neanderthal ancestry remains in some AMHs. We consider genomic regions in Europeans identified by the Sprime algorithm that contain a high density of single nucleotide variants absent in unadmixed African populations and that frequently match the Altai Neanderthal allele.^34^ Filtering for high confidence Neanderthal introgressed regions, we identified 1345 segments of the human genome tolerant of Neanderthal ancestry that have a median length of 299 kb (IQR: 174 – 574 kb), covering 19% of the genome (Methods, Fig. S1A). These Neanderthal ancestry segments have, on average, 116 putative introgressed variants comparable to the Altai genome, with 76% of them matching the Altai allele (Fig.S1B-C). However, as expected, most variants segregating in Eurasian populations in these regions of the genome that tolerate Neanderthal ancestry are not of Neanderthal origin (Fig. 1A).

**Figure 1.**
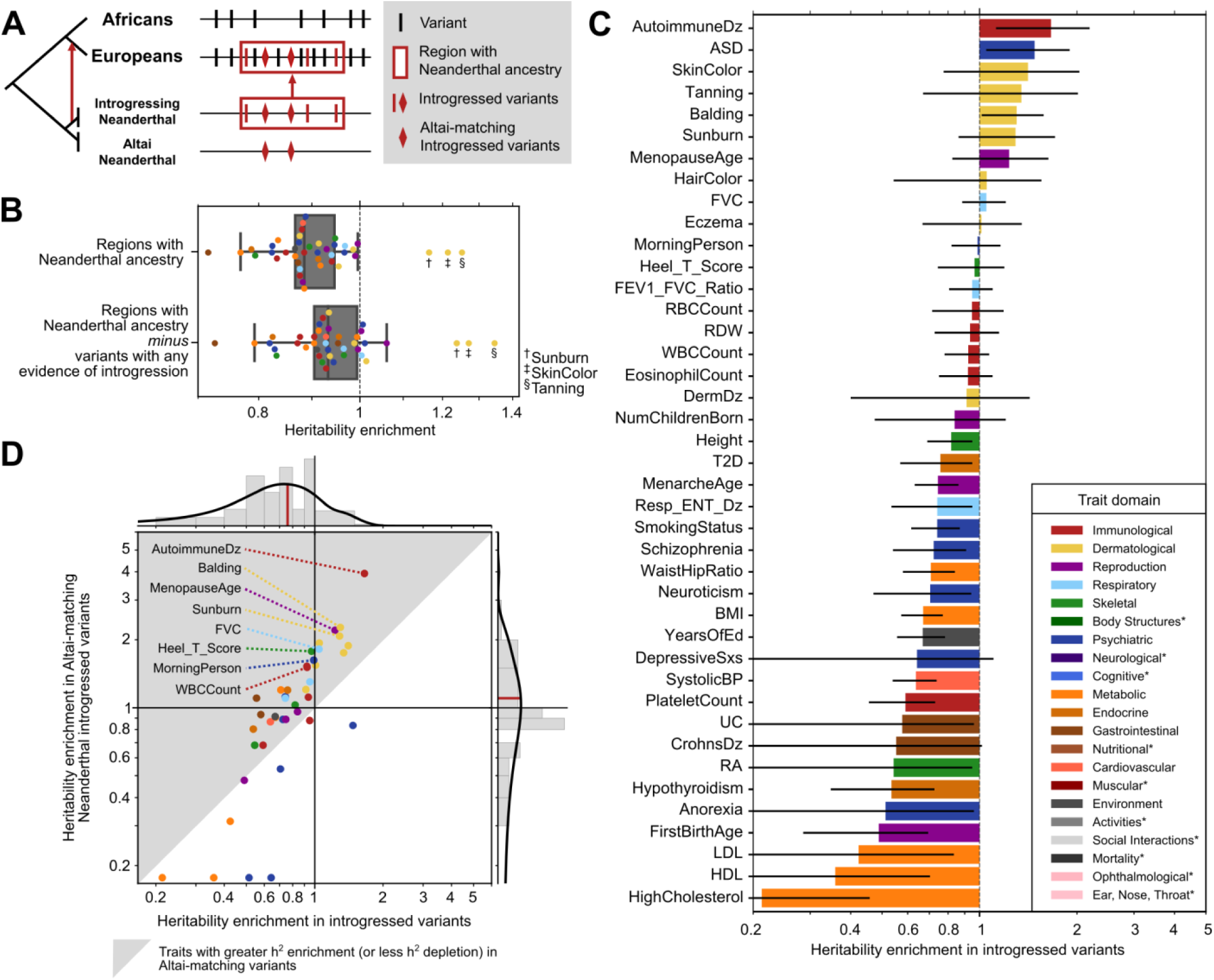
Complex trait heritability is broadly depleted in introgressed regions and variants, yet introgressed variants have an outsized effect on certain traits. (A) Segregating variants (vertical lines and diamonds) considered in this study. We focus on variants inside regions of the human genome tolerant of Neanderthal ancestry (red box). These alleles have multiple evolutionary histories: most are segregating on non-introgressed haplotypes (black), many are present in Eurasians due to introgression (red), and some of the introgressed alleles were shared among multiple Neanderthal populations including both the Altai Neanderthal and the introgressing Neanderthal population (diamonds). (B) Regions of the genome where Neanderthal ancestry remains are depleted for heritability of 41 diverse complex traits (0.88x background expectation, *P =* 8×10^−7^) except for sunburn, skin color, and tanning. Removing introgressed variants (LD expanded to *r*^2^ > 0.5), these regions are still broadly depleted for trait heritability (0.93x, *P =* 0.003). Each dot represents the heritability enrichment or depletion for a single trait in regions of the genome with Neanderthal ancestry. (C) Most (76%) traits trend toward depletion in specific variants with any evidence of introgression. Error bars represent standard errors. Traits are colored by their domain (legend); starred domains appear only in later figures. These colors will be used in all figures. (D) Compared to trait heritability in variants with any evidence of introgression (x-axis, median: 0.76x), Altai-matching introgressed Neanderthal variants (y-axis, median: 1.10x) are more enriched for heritability for 78% of traits (gray triangle). Traits that trend towards heritability enrichment include immunological, dermatological, lung capacity, bone density, chronotype, and menopause-related traits. Traits with depletion less than 0.125 are truncated.

To estimate heritability enrichment or depletion in these regions where Neanderthal ancestry remains, we conducted partitioned heritability analysis using stratified-LD score regression (S-LDSC).^35,36^ In this context, heritability depletion indicates that variation in regions with Neanderthal ancestry is less associated with phenotypic variation than expected given a null hypothesis of complete polygenicity; heritability enrichment means that the variants associate with more of the phenotypic variation than expected. To start, we considered summary statistics from a previously-curated representative set of 41 diseases and complex traits with high-quality GWAS (average number of individuals [N] = 329,378; SNPs in GWAS [M] = 1,155,239; h^2^_SNP_ = 0.19; Table S1).^37–48^

Regions tolerant of Neanderthal ancestry are broadly depleted of complex trait heritability (Fig. 1B). Across traits, these regions are 1.13-fold depleted for the heritability expected from the background genome (*P* = 8×10^−7^). After removing variants with evidence of introgression in any Eurasian populations (LD expanded to *r*^2^ > 0.5 [Methods]), these regions are still 1.07-fold depleted for trait heritability (*P =* 0.003). The relative increase in the heritability enrichment after removing introgressed variants (and those in high LD with them) suggests that introgressed variants account for much, but not all, of the heritability depletion in these regions tolerant of Neanderthal ancestry. The depletion across traits also holds for introgressed haplotypes identified by the earlier S* method (Supplemental Fig. S2A).^49^ These analyses demonstrate that regions of the genome that retain Neanderthal ancestry not only have less evidence for constraint and function at the molecular level,^25–27^ but they are also depleted for variation influencing a diverse array of complex traits.

We find three exceptions to this complex trait heritability depletion; sunburn (1.16x, *P* = 0.3), skin color (1.21x, *P* = 0.4), and tanning (1.25x, *P* = 0.4) are not depleted among regions that retain Neanderthal ancestry (Fig. 1B). These three traits are genetically correlated with magnitudes between *r* = 0.55 and 0.86. Several previous hypotheses suggest that the introgression of Neanderthal alleles related to hair and skin pigmentation could have provided non-African AMHs with adaptive benefits as they moved to higher latitudes.^3,4,17,18^ The exceptions to the overall depletion we observe for skin traits suggest that regions of the human genome involved in skin pigmentation were more tolerant of Neanderthal introgression than regions of the genome associated with other traits.

### Neanderthal introgressed variants are broadly depleted for complex trait heritability

Genomic regions with remaining Neanderthal ancestry in AMH contain many introgressed variants; however, the majority of variants segregating in these regions are not of Neanderthal origin and are not exclusive to introgressed haplotypes (Fig. 1A). We will refer to such variants as “non-introgressed” for simplicity. In the previous section, we demonstrated that non-introgressed variants in regions with remaining Neanderthal ancestry are generally depleted for heritability of most complex traits. In this section, we focus on the heritability in introgressed variants specifically. Introgressed variants have diverse evolutionary origins and histories, and these histories suggest different expectations for their functional effects. For example, older introgressed alleles that segregated in the Neanderthal lineage for hundreds of thousands of years might be more tolerated than younger introgressed alleles specific to the introgressing Neanderthal group, because they have been maintained by selection in more diverse populations and environments.

Thus, we quantified the relationship between heritability of the representative 41 complex traits and four sets of Neanderthal-introgressed variants identified by Sprime.^34^ The largest set included all variants with evidence of introgression in any Eurasian population (N = 900,902, Methods); this set will be referred to as “introgressed variants” throughout the manuscript. This set includes not only high-confidence Neanderthal-origin introgressed variants, but also some ancestral alleles lost in Africans that were reintroduced to Eurasians through archaic introgression^5^, variants with origins in other archaic hominins such as Denisovans, and variants tightly linked to introgressed haplotypes that were introduced in Eurasians shortly after introgression. The most stringent set includes Neanderthal-introgressed alleles that explicitly match the Altai genome and are observed in Europeans (N=138,774, Methods); this set will be referred to as “Altai-matching introgressed variants.” This set represents a high-confidence set of variants of Neanderthal origin that were likely common among diverse Neanderthal groups given the substantial divergence of the Altai Neanderthal from the introgressing population.^1,50^ However, it excludes many true introgressed Neanderthal alleles, such as those that are not present in the Altai Neanderthal. We calculated partitioned heritability on these two sets and two other intermediate-stringency sets; results from all sets are in Fig. S3.

Consistent with our observations in regions tolerant of introgression (Fig. 1B), 31 of the 41 (76%) traits trend towards depletion of heritability in the set of introgressed variants (Fig. 1C). We observed significant depletion for heritability of cholesterol level (4.6-fold depletion, Benjamini-Hochberg FDR-corrected, *q* = 0.02), platelet count (1.7-fold depletion, *q* = 0.04), systolic blood pressure (1.6-fold depletion, *q =* 0.01), years of education (1.5-fold depletion, *q* = 0.04), and body mass index (BMI, 1.5-fold depletion, *q* = 0.02). Next, we calculated partitioned heritability for the Altai-matching introgressed variants. Despite the overall depletion for complex trait heritability in regions of the genome tolerant of introgression and all introgressed variants, these Altai-matching variants trend towards heritability depletion in only 19 of 41 (46%) traits (Fig. 1D, Fig. S3A). The median trait heritability in the Altai-matching variants is 1.10-fold *enriched*, compared to the 1.32-fold *depletion* in the broader set of introgressed variants (Fig. 1D) and the 1.13-fold *depletion* in regions tolerant of Neanderthal ancestry (Fig. 1B).

The traits that trend towards heritability enrichment in Altai-matching Neanderthal-introgressed variants include immunological (autoimmune disease, white blood cell [WBC] count), hair (balding, hair color), skin (sunburn, tanning, skin color), lung capacity (FVC), bone density (Heel T-Score), chronotype (morning person) and menopause (menopause age) related traits (Fig. 1C-D, *P =* 0.02-0.29, Fig. S3). We note that most do not meet strict multiple testing corrected p-value thresholds for enrichment; however, the results are highly correlated (R^2^ = 0.79) with partitioned heritability estimates for introgressed variants replicated using the S* approach (Fig. S2B-C).^49^

Overall, we observed several trends in the enrichment patterns among the 41 traits considered in the Neanderthal introgressed variant partitioned heritability analysis. The majority of dermatological (7/7), immunological (4/5), and respiratory (3/3) traits trended towards enrichment in the Altai-matching introgressed variants, while metabolic (4/5) and psychiatric (5/7) traits trended toward depletion (Fig. 1D). However, these traits represent a small subset of the available GWAS across diverse trait domains.

### Over 405 complex traits, Neanderthal introgressed variants are most enriched for heritability of dermatologic traits and depleted for cognitive traits

To expand our analysis and evaluate trends across a broader set of traits, we analyzed GWAS summary statistics from the UK BioBank and FinnGen.^37,51,52^ Following guidelines for S-LDSC analyses^36,51^, we only considered GWAS for traits that: (1) met criteria for high confidence estimates of SNP heritability (N_eff_ *>* 40,000, low SE, low sex bias [Methods]) and (2) were significantly heritable (z > 7, P < 1.28×10^−12^). This identified 405 GWAS (average N = 288,130, h^2^_SNP_ = 0.108, Table S2). We assigned these GWAS to the domains, chapters, and subchapters from the GWAS Atlas.^53^ For traits unclassified in the GWAS Atlas, we assigned hierarchical classifications manually. We performed partitioned heritability analysis on these traits using all sets of Neanderthal-introgressed variants described above (Figs. S4-S6); here, we report results from the most stringent Altai-matching introgressed variants (Fig. 2).

**Figure 2.**
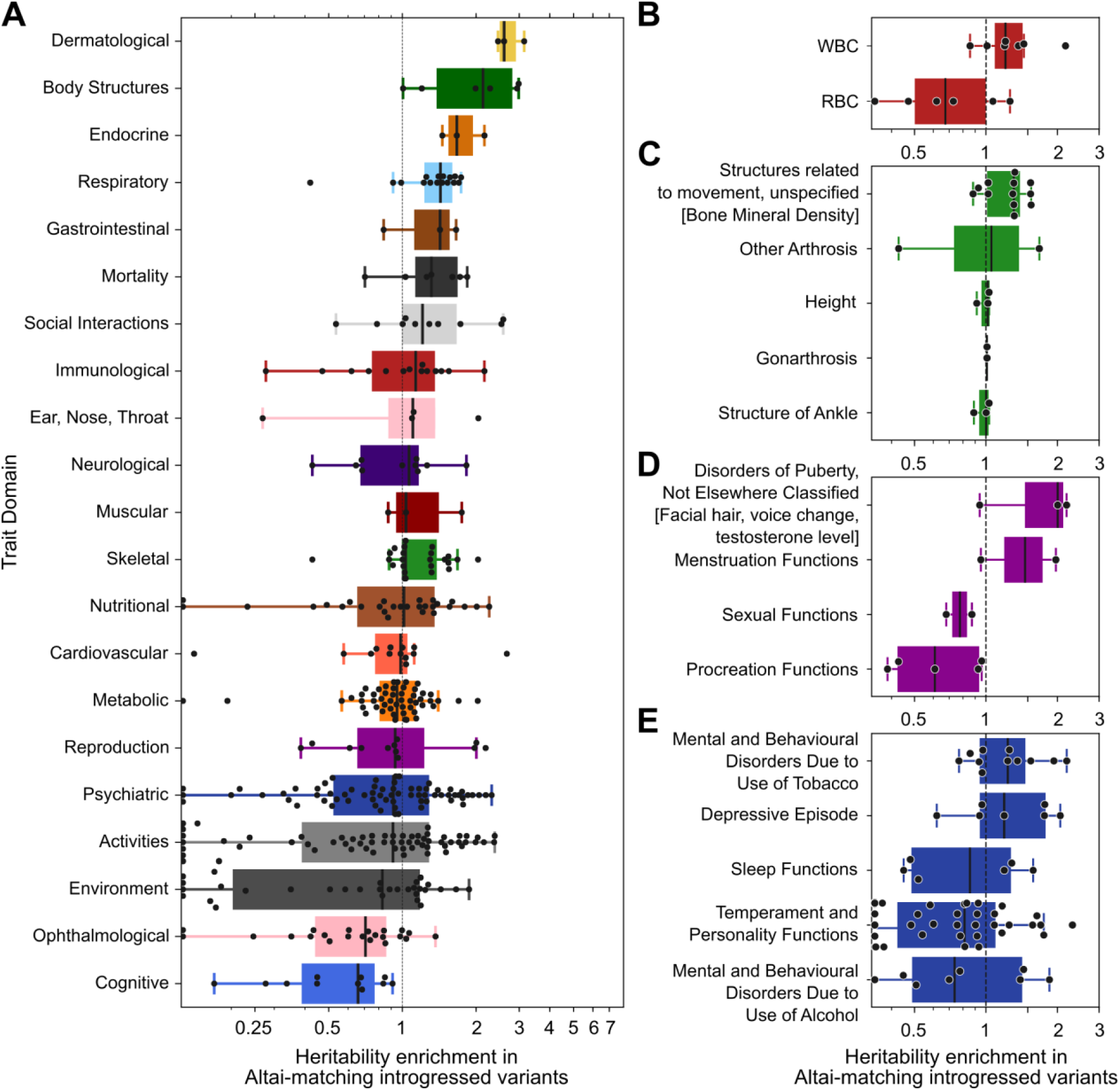
Heritability enrichment and depletion in introgressed variants across 405 traits clustered by domain. (A) Introgressed variants are most enriched for dermatological-domain traits (hair-related) and most depleted for cognitive-domain traits (higher-level cognitive and memory functions). Each point represents heritability enrichment or depletion of one trait in Altai-matching introgressed variants. Traits with depletion less than 0.125 are plotted on the baseline for visualization. Within some domains, introgressed variants also show variable heritability enrichment. We quantified enrichment and depletion for traits at the more granular chapter and subchapter levels. (B) Dividing immunologic traits into subchapters, WBC-related traits trend toward heritability enrichment (1.2-fold enriched, *P =* 0.07) in Neanderthal variants, while RBC-related traits trend towards depletion (1.5-fold depletion, *P =* 0.12). (C) For skeletal traits, bone mineral density-related traits show the most enrichment for heritability in introgressed variants (1.3-fold enrichment, *P =* 0.004). (D) For reproductive traits, puberty- and menstruation-related traits are nominally enriched for heritability (2.0-fold enrichment, *P =* 0.1), whereas sexual and procreation functions are depleted (1.5-fold depleted, *P =* 0.02). (E) For psychiatric traits, tobacco use disorders (1.2-fold enrichment, *P =* 0.09) and depression-related traits (1.2-fold enrichment, *P =* 0.25) trend towards enrichment. Domain, chapter, and subchapter-level results across all traits are similar using multiple sets of introgressed variants (Figs. S4-S6, Table S3-4).

In this diverse set of traits, Altai-matching introgressed variants are most enriched for heritability of dermatologic (hair-related) traits (2.6-fold enriched, *P =* 0.006, one sample T-test) and most depleted for cognitive (higher-level cognitive and memory functions) traits (1.5-fold depleted, *P* = 0.002) (Fig. 2A, Table S3). Body structure (*e.g.* fractures, dental diseases, 2.1-fold enriched, *P* = 0.02), endocrine (1.7-fold enriched, *P =* 0.04), and respiratory (1.4-fold enriched, *P =* 0.01) traits also exhibit strong trends towards enrichment. Traits related to eye structure (1.4-fold depleted, *P =* 0.03), daily activities (1.1-fold depleted, *P =* 0.001) and environment (*e.g.* jobs, finances, 1.2-fold depleted, *P =* 0.01) are depleted in addition to cognitive traits. The depletion in cognitive traits demonstrates that the strong depletion for Neanderthal alleles in regulatory regions active in the brain influenced relevant complex traits.^5,26,54^

Other trait domains exhibit substantial intra-domain diversity in the heritability patterns with some traits showing strong enrichment and others depletion in Altai-matching introgressed variants. Thus, we also quantified enrichment and depletion for traits at the more granular chapter and subchapter levels. Dividing immunologic traits into subchapters, WBC-related traits trend toward heritability enrichment (1.2-fold enriched, *P =* 0.07) in Altai-matching variants, while RBC-related traits trend towards depletion (1.5-fold depletion, *P =* 0.12, Fig. 2B). For skeletal traits, bone mineral density-related traits show the most enrichment for heritability in introgressed variants (1.3-fold enrichment, *P =* 0.004, Fig. 2C). For reproductive traits, puberty- and menstruation-related traits are enriched for heritability (2.0-fold enrichment, *P =* 0.1), whereas sexual and procreation functions are depleted (1.5-fold depleted, *P =* 0.02), Fig. 2D), possibly reflecting barriers to introgression. For psychiatric traits, tobacco use disorders (1.2-fold enrichment, *P =* 0.09) and depression-related traits (1.2-fold enrichment, *P =* 0.25, Fig. 2E) trend towards enrichment, consistent with previous observations.^3,4^ Domain, chapter, and subchapter-level results across all traits for all the sets of introgressed variants are in Figs S4-S6 and Tables S3-S4.

### Neanderthal alleles confer directional effects for some traits

Partitioned heritability analyses quantify the overall contribution of introgressed loci to variation in traits across humans; however, they do not test for directionality of the effects of the introgressed alleles on traits. We now test whether introgressed alleles consistently have effects in the same direction (*e.g.*, risk increasing) for eight traits spanning diverse phenotypic domains that were enriched for partitioned heritability in Altai-matching introgressed variants (AutoimmuneDz, Balding, Sunburn, FVC, Heel_T_Score, MorningPerson, MenopauseAge, WBCCount, Fig. 1C). We quantify Neanderthal introgressed allele direction of effect in two ways.

First, focusing on the trait-associated variants with strongest effects, we intersected Altai-matching introgressed alleles with genome-wide significant variants from the eight GWAS. We then quantified if there is overrepresentation of introgressed alleles in the risk-increasing or risk-decreasing direction. We considered GWAS variants significant at *P* < 1 × 10^−8^ and pruned variants in perfect LD (*r*^2^ = 1) to reduce redundant counts due to linked loci. Results from using less strict thresholds (*P* < 5 × 10^−8^, *P* < 1 × 10^−6^ and *r*^2^ > 0.8, *r*^2^ > 0.5) are similar (Fig. S7).

Four traits show a significant difference (*P* < 0.05, χ^2^ goodness of fit test) in the direction of effect of introgressed genome-wide significant variants: balding, menopause age, forced vital capacity, and morning person (Fig. 3A). For balding, the Altai-matching introgressed alleles were 7.5 times more likely to be significantly associated with hair loss (less Type 1 Balding, *P* = 0.002). For menopause age, the introgressed alleles were 5 times more associated with younger age at menopause (*P* = 0.02). For forced vital capacity, the introgressed alleles were 2.5 times more associated with larger lung volumes (*P =* 0.01). For morning person, the introgressed alleles were 4.3 times more associated with increased likelihood of being a morning person, as compared to evening person (*P* = 0.01). Additionally, introgressed alleles associated with Heel T Score and Sunburn traits have non-significant directional trends that support previous findings. More of the associated introgressed alleles are associated with increased bone density (1.4x, *P =* 0.12) and with increased sunburn risk (4x, *P* = 0.18).

**Figure 3.**
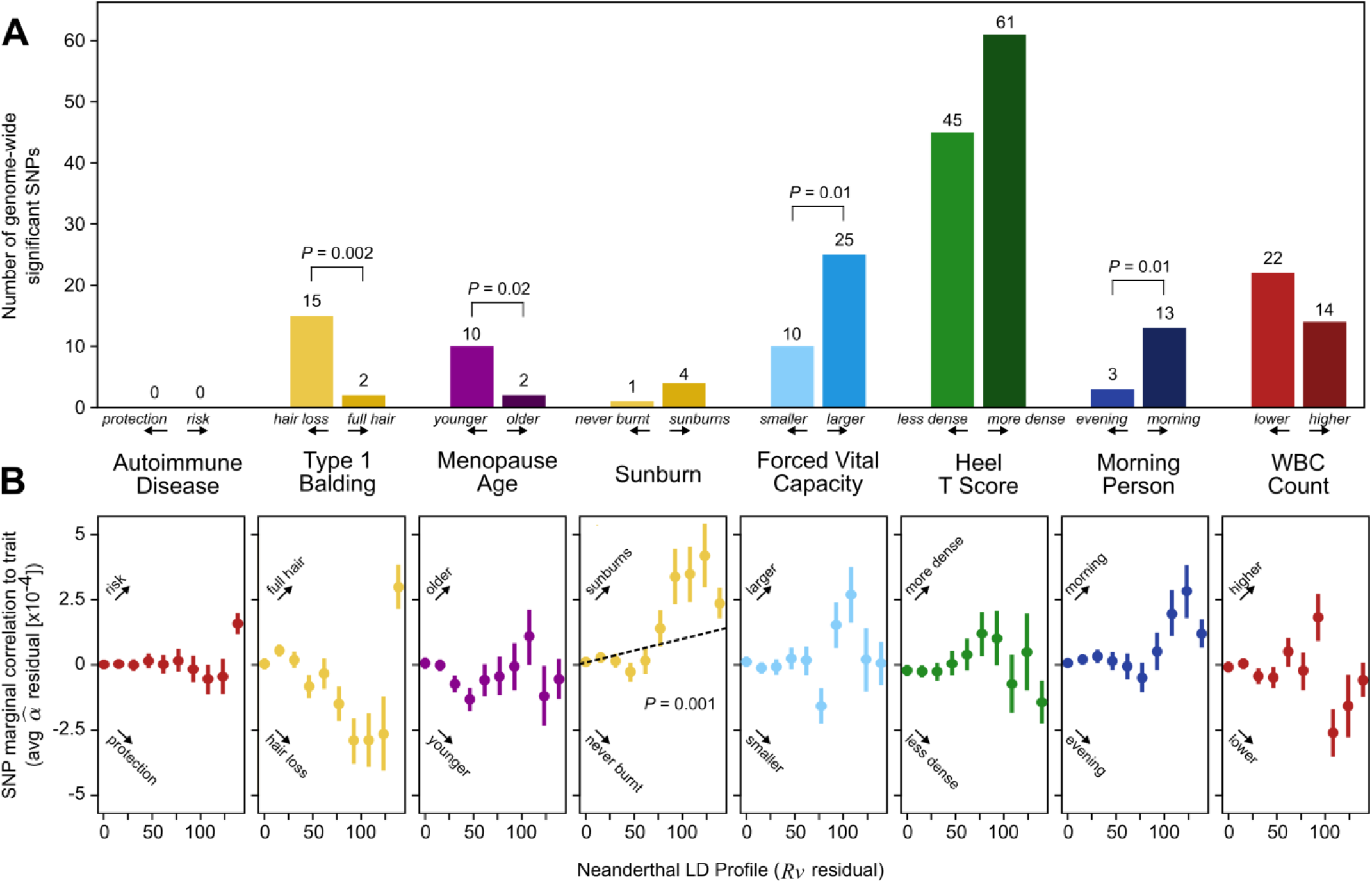
Neanderthal alleles confer directional effects for some traits. For eight traits with partitioned heritability enrichment in Neanderthal introgressed variants (Fig. 1D), we assessed the direction of effect of the Neanderthal alleles with two approaches. (A) The first intersects introgressed Altai-matching Neanderthal alleles (LD-expanded to *r*^2^ = 1) with the genome-wide significant (*P* < 1 × 10^−8^) variants from each GWAS. For each trait, we plot the number of significant variants (variants in perfect LD [*r*^2^ = 1] are pruned) by the direction of effect of the Neanderthal allele. Four traits show a significant difference (*P* < 0.05, χ^2^ goodness of fit test) in direction of effect: increased balding, younger menopause age, increased forced vital capacity, and morning person. For example, of the 17 Neanderthal alleles associated with balding, 15 are associated with hair loss and only two with full hair. (B) The second method uses signed LD profile regression to consider the direction of effect over all variants, not just those with genome-wide significant effects. For each variant, we plot the marginal correlation (α^) of the variant to the trait versus the Neanderthal LD profile (*R*ν). Increased signed LD Profile reflects increased conditional LD to Neanderthal introgressed allele. For visualization, we bin *R*ν into 10 equally spaced intervals and plot the average α^ with 95% bootstrapped confidence intervals. The correlation between *R*ν and α^ indicates the genome-wide direction of effect. For example, the positive correlation for sunburn indicates a significant relationship genome-wide between Neanderthal introgressed alleles and risk for sunburn (*P* = 0.001).

By considering only genome-wide significant variants, this directionality analysis tests for a relationship at the extreme of effect size and significance. To assess if there is directional bias among Neanderthal-introgressed alleles on these eight traits across the effect distribution, we used signed LD profile regression.^55^ Signed LD profile regression assesses whether variant effects on a trait (from GWAS summary statistics) are correlated genome-wide with a signed genomic annotation via the *signed LD profile* (*R*ν). Our genomic annotation is a vector that quantifies LD to an Altai-matching Neanderthal introgressed allele. Per variant, a higher *Neanderthal LD profile* (*R*ν) means that the variant aggregately tags variants with higher LD to introgressed allele(s). Per variant, a higher marginal effect (α^) means that the allele is correlated in a positive direction to the trait (relative to the coding of the GWAS). Therefore, a genome-wide positive correlation between α^ and *R*ν indicates a positive relationship between LD to introgressed allele(s) and increase in the trait. Conversely, a genome-wide negative correlation between α^ and *R*ν indicates a negative relationship between LD to introgressed allele(s) and a decrease in the trait.

Using signed LD profile regression, we find a significant genome-wide correlation between higher LD to introgressed alleles (Neanderthal LD profile, *R*ν) and increased risk for Sunburn α^ (*r*_*f*_ *=* 0.18%, *P =* 0.001, Fig. 3B, Table S5). Other traits show directionality similar to the significant hit analysis that does not reach significance when considering the entire genome. For example, Neanderthal LD profile correlates with risk for younger menopause age and increased propensity to be a morning person. The remaining traits do not show a consistent directionality genome-wide; instead, these traits have genomic windows where Neanderthal alleles contribute significantly in risk-increasing and other windows with risk-reducing directions. Analysis of all 41 traits considered in Fig. 1 demonstrated that introgressed alleles do not confer directional effects for most traits. However, remaining introgressed alleles do have significant genome-wide correlations with protection from anorexia (*r*_*f*_ = -0.93%, *P*<1×10^−6^) and schizophrenia (*r*_*f*_ *= -0.27%, P* = 6×10^−4^; Table S5).

### Identifying introgressed Neanderthal alleles with effects on human traits

The above analyses identify several traits for which variants of Neanderthal ancestry have an outsized effect. In this section, we use these results to identify illustrative examples of how specific introgressed alleles may influence these traits. In contrast to previous studies that simply intersected introgressed alleles with effects on traits from association studies, we locate regions of interest based on strong correlations between LD to Altai-matching introgressed alleles (*Neanderthal LD profile, R*ν) and trait-associated risk or protection (α^) in sliding windows across the genome (Methods).

We first considered the loci most strongly contributing to the genome-wide significant positive correlation between Neanderthal LD profile and sunburn risk. One such window with a strong positive correlation (chr9:16641651–16787775, R = +0.83) overlapped the gene *BNC2* which influences skin pigmentation levels in Europeans (Fig. S8).^56^ As previously reported, *BNC2* has strong support for the association of introgressed variation with risk for sunburn: it is overlapped by a Neanderthal haplotype at ∼70% frequency in Europeans associated with sun sensitivity^17,57^ and is near archaic alleles that tag introgressed haplotypes (rs10962612 tagging chr9:16,720,122–16,804,167; rs62543578 tagging chr9:16,891,561–16,915,874) strongly associated with increased incidence of childhood sunburn and poor tanning in the UK Biobank.^4^ We report 29 other windows associated with sunburn using this method in Table S6. Among these regions, we identify many promising candidates, including one nearby *SPATA33* which has been implicated in tanning response, facial pigmentation, and skin cancer^58^ and one nearby *MC1R* which is a key genetic determinant of pigmentation and hair color.^4^

We apply this method to the eight other traits and report windows of interest in Table S6 including windows recovering previous links between Neanderthal introgression and chronotype surrounding *ASB1* (overall R = -0.92, Fig. S8).^4^ Recapitulating these established findings highlights the utility of this method for identifying regions where Neanderthal introgression may affect modern Europeans.

We identified a novel relationship between Neanderthal alleles and morning person status near *NMUR2*. Two windows within chr5:151745423-151931514 show a positive relationship between Neanderthal LD profile and morning person status (overall R = +0.91, Fig. 4A). This positive correlation suggests that increased LD to Neanderthal alleles is associated with increased propensity to be a morning person. In addition to this window being physically near *NMUR2*, 168 of 169 introgressed variants that overlap this window are expression quantitative trait loci (eQTL) in GTEx^59^ negatively regulating *NMUR2* in frontal cortex cells (as expected, these variants are in high LD; for tag variant rs4958561: *P =* 1×10^−9^, Fig 4A). For illustration purposes, we pick the introgressed variant with the highest α^ (rs4958561) to tag the Neanderthal introgressed variation in this region. However, we recognize that this region’s association with the Morning Person GWAS and *NMUR2* expression is likely from other linked variants. In a PheWAS, the top ten traits most associated with rs4958561 include ease of getting up in the morning (N = 2, *P =* 1×10^−14^ & 2×10^−14^), chronotype (N = 2, 2×10^−12^ & 4×10^−12^), morningness (N = 2, *P =* 4×10^−12^ & 2×10^−11^), sedentary behavior (*P =* 4×10^−9^), and tea intake (*P =* 1×10^−8^) (all pass Bonferroni correction for testing 4756 GWAS).^53^ *NMUR2* encodes the Neuromedin-U receptor 2, which is acted on by neuromedins U (NMU) and S (NMS) to phase shift circadian activity rhythm.^60–62^ *In vivo*, NMU shows circadian expression in rat brains in response to melatonin and a genetic overexpression screen in zebrafish larvae identified *Nmu* to promote hyperactivity through Nmu receptor 2.^63,64^ Ultimately, we hypothesize that Neanderthal introgressed alleles or tightly linked alleles downregulate *NMUR2* in the brain leading to an association with increased morning person propensity.

**Figure 4.**
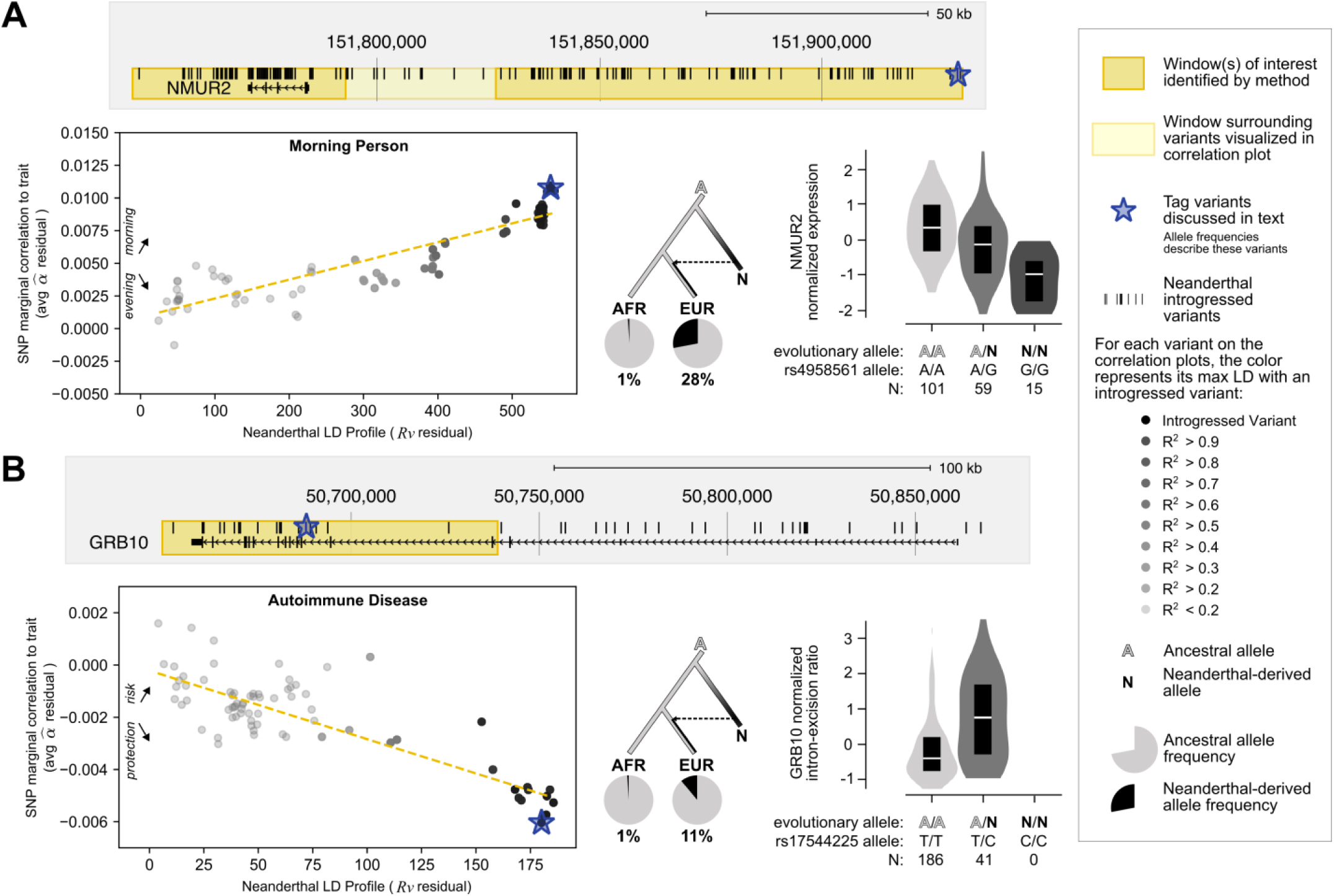
Windows with strong correlations between Neanderthal LD profile and trait-association highlights putative mechanisms for the effect of introgression on chronotype and autoimmunity. (A) The genomic region shown highlights the novel positive relationship between Neanderthal LD profile and morning person status in two regions (chr5:151,745,423-151,793,214, chr5:151,826,774-151,931,514) overlapping and nearby *NMUR2*. The scatter plot shows chr5:151,745,423-151,931,514’s positive relationship (R = +0.91). The starred variant (rs4958561) is an eQTL in which Neanderthal alleles increase *NMUR2* expression in frontal cortex (*P* = 1×10^−9^) and cortex (not shown; *P* = 9×10^−5^). (B) The genomic region shown highlights an identified window (chr7:50,649,920-50,739,129) overlapping *GRB10* which has a negative relationship (R = -0.84) between Neanderthal LD profile and autoimmune disease risk (hence, increased Neanderthal LD profile is associated with protection from autoimmunity). This relationship would not have been identified by intersecting introgressed alleles with genome-wide significant GWAS associations. The starred variant (rs17544225) is an sQTL in which Neanderthal alleles increase *GRB10*’s intron excision in spleen (*P* = 3×10^−9^). For each example, we display the genomic region overlapped by the identified window(s) of interest (dark yellow box), genes, all Altai-matching Neanderthal-introgressed variants (black marks). For the region in light yellow, we display a scatter plot between the variant’s Neanderthal LD Profile (*R*ν) and trait marginal correlation (α^). Each variant is colored by its maximum LD to an introgressed variant. We display the evolutionary history (dendrogram) of each discussed tag variant (blue star) with its allele frequency in the in the 1000G African (AFR) and Europeans (EUR) super populations (pie charts).

Our directional correlation-based method provides stronger evidence for a causal relationship with Neanderthal variants than an intersection between introgressed alleles and genome-wide significant variants because directional effects are less confounded by genomic co-localization of Neanderthal ancestry with other functional elements.^55^ Furthermore, with this method we can identify trait-associated regions that are not explicitly tagged by a genome-wide significant introgressed allele, yet still have a significant directional relationship between Neanderthal LD profile and a trait. For example, there were no Altai-matching introgressed alleles had genome-wide significant associations with autoimmune disease. However, we identify a window with a strong negative correlation indicating a protective relationship between Neanderthal introgression and autoimmune disease (chr7:50649920-50739129, R = -0.84, Fig. 4B). In this ∼90 kb window, there are six GWAS tag variants that overlap Neanderthal alleles; these variants are nominally associated with autoimmune disease protection (*P* = 5×10^−5^ - 5×10^−4^). However, in this region, there are 55 additional unique GWAS tag variants in LD (*r*^2^ > 0.2) with these six Neanderthal alleles that provide power to test if the GWAS signal in this region is likely related to the Neanderthal alleles versus other nearby variation.

This region with a protective relationship between Neanderthal LD profile and autoimmune disease protection flanks *GRB10*, which encodes growth factor receptor-bound protein 10 known to interact with a number of tyrosine kinase receptors and signaling molecules.^65^ Although *GRB10’*s role in immunity is still unclear, it has been associated with a subtype of systemic sclerosis (lcSSc); patients with systemic sclerosis have higher expression of *GRB10* in monocytes.^66,67^ Studies of *Grb10* deficient mice demonstrated *Grb10*’s role in hematopoietic regeneration *in vivo*.^68^ Additionally, in a transcriptome study of CD4+ Effector Memory T cells (CD4^+^ T_EM_), *GRB10* was the most significantly downregulated gene after T-cell receptor stimulation.^69^ Notably, in both human and mouse, *GRB10* mRNA is highly alternatively spliced, resulting in four to seven unique isoforms.^70^ Of the 20 introgressed variants overlapping this window, 17 are splicing quantitative trait loci (sQTL, increasing intron excision, for tag variant rs17544225: *P* = 3×10^−9^, Fig 4B), in the spleen.^59^ Again, we pick we pick the introgressed variant with the highest α ^ (rs17544225) to tag the Neanderthal introgressed variation in this region. In a PheWAS, the top ten traits associated with rs17544225 include monocyte count (N = 2, *P =* 4×10^−8^ & 3×10^−7^) and monocyte percentage (N = 2, *P =* 3×10^−6^ & 4×10^−6^).^53^ Therefore, we hypothesize that Neanderthal introgressed alleles or their tightly linked alleles play a role in regulation or splicing of *GRB10* contributing to changes in monocytes that may lead to protection from autoimmunity.

## DISCUSSION

Here we apply powerful methods to estimate the heritability patterns across more than 400 diverse traits in genomic regions influenced by Neanderthal introgression. Regions tolerant of Neanderthal ancestry in modern populations are depleted of heritability for all traits considered, except those related to skin and hair. Introgressed alleles are also depleted for heritability of most traits; however, there is modest enrichment for heritability of several traits among alleles with high-confidence Neanderthal origins, including autoimmune disorders, hair and skin traits, chronotype, bone density, lung capacity, and age at menopause (Fig. 1, 2C). Summarizing these heritability patterns over trait domains, we find that dermatological, endocrine, and respiratory traits are consistently enriched for heritability among Altai-matching Neanderthal introgressed variants, whereas cognitive and ophthalmological domains are the most depleted. Additionally, several trait domains show divergent heritability patterns, *e.g.* among psychiatric and reproductive traits (Fig. 2D-E). Using two methods for evaluating the direction of effect of variants on traits, we find significant directional biases for introgressed alleles with balding risk, younger menopause age, sunburn risk, forced vital capacity increase, and morning preference (Fig. 3). Finally, we show how our new approaches can highlight novel candidate introgressed variants that impact risk for disease (Fig. 4).

Our results expand the current understanding of the functional effects of introgressed variants in several dimensions. First, previous studies of regions tolerant of Neanderthal ancestry have found depletion for evidence of background selection and functional annotations, such as genes and gene regulatory elements active in specific tissues.^17,18,25,27^ We move beyond these proxies for function and show depletion for effects on diverse complex traits in a human population. This further suggests selection against Neanderthal introgression in these trait-associated genomic regions. However, we also find a striking exception to this pattern for variation associated with skin color and tanning, which is consistent with previous hypotheses that genomic regions associated with skin traits tolerated adaptive introgression.^3,18^ This also agrees with previous attempts that tested for genome-wide effects of Neanderthal ancestry on complex traits which also found enrichment for traits related to skin and hair. However, our method expands on these results by agnostically testing over 400 traits across many domains rather than selecting 46 specific traits as in GCTA analysis in Simonti *et al.* 2016.^3^ Furthermore, our partitioned heritability method for identifying enrichment considers the full distribution of variant effect sizes from the GWAS rather than selecting an *ad hoc* significance threshold (*P* < 1.0 × 10^−8^) as in the analysis of 136 traits by Dannemann *et al.* 2017.^4^ As a result, our analysis of over 400 traits using S-LDSC provides greater power and range to identify risk variance explained by Neanderthal variation.

Second, we separately analyze trait heritability patterns in regions tolerant of Neanderthal ancestry, in remaining introgressed variants, and in non-introgressed variants (Fig. 1A). Considering these variant sets separately enables us to interpret of the effects of introgressed regions in their genomic context. For example, we find modest enrichment for heritability of several traits among introgressed alleles, even though they are in regions of the genome with overall depletion for heritability of these traits. Our analyses also suggest differences in heritability among different subsets of introgressed variants. The introgressed variants that remain in AMH genomes are the result of complex selective and demographic pressures following admixture.^8,23,25^ Introgressed haplotypes carry alleles of different origins, including ancestral alleles lost in some modern Eurasian populations, and in some cases, these non-Neanderthal variants are responsible for the functions of introgressed haplotypes.^5^ Supporting this, our analysis of different sets of alleles on introgressed haplotypes revealed that introgressing alleles matching the Altai Neanderthal are less depleted for heritability than other introgressed alleles. The introgressing Neanderthal population diverged from the Altai Neanderthal population more than 100 kya.^1,50^ Given this substantial divergence, we anticipate that the Altai-matching introgressed alleles were more tolerated and likely at higher frequency in different Neanderthal populations. This suggests that Altai-matching introgressed alleles would be less likely to have strong deleterious effects than other younger introgressed Neanderthal alleles, potentially explaining the lower levels of depletion (and modest enrichment for some traits) of heritability in these variants.

Third, we introduce a new approach for testing for consistent direction of effects for introgressed alleles on traits. Using this approach, we show that Neanderthal introgression generally increased propensity for sunburn, balding, larger lung capacity, and younger menopause, while it had both increasing and decreasing effects on most other traits. With this directionality metric, we highlight candidate functional introgressed variants involved in autoimmune disease that would not have been identified by simply intersecting introgressed alleles with GWAS results.

Together, the patterns of heritability and direction of effect reveal contrasting patterns of selection since admixture on introgressed variation associated with different traits (Fig. 5A). Along the dimensions of heritability enrichment vs. depletion and directional bias vs. no bias, traits fall into four general quadrants (Fig. 5B). First, most traits show heritability depletion among introgressed variants and no bias in the direction of effect. This suggests selection against introgressed variants that influenced these traits (Fig. 5B, bottom left). Second, the opposite pattern—enrichment for heritability in introgressed variants and a bias in their direction of effect—suggests that introgression introduced functional alleles that were positively selected in AMHs (Fig. 5B, top right). For example, the enrichment for heritability of sunburn and tanning in introgressed alleles and the bias in direction of effect in AMH suggests that these introgressed alleles decreased hair and skin protection against sun exposure in ways that may have been beneficial, perhaps in response to decreased UV at higher latitudes. Third, traits, like autoimmune disease risk and WBC count, have heritability enrichment among introgressed variants, but no directional bias. In this case, introgression likely contributed increased diversity—both trait increasing and decreasing—relevant to the trait into AMHs that was beneficial as they adapted to non-African environments (Fig. 5B, bottom right). This further suggests that Neanderthals themselves varied in these traits.^4^ Fourth, traits like anorexia and schizophrenia, show depletion for heritability among introgressed variants, but in contrast to most depleted traits, they have a significant directionality bias in the effects of remaining introgressed variants (Fig. 5B, top left). We hypothesize that this pattern could be produced by negative selection purging most introgressed alleles that influence the trait paired with selection for a small number of introgressed protective alleles. Supporting this interpretation, remaining Altai-matching variation has the most significant correlation with protective benefit against serious fitness-reducing diseases (anorexia, schizophrenia).^71^

**Figure 5.**
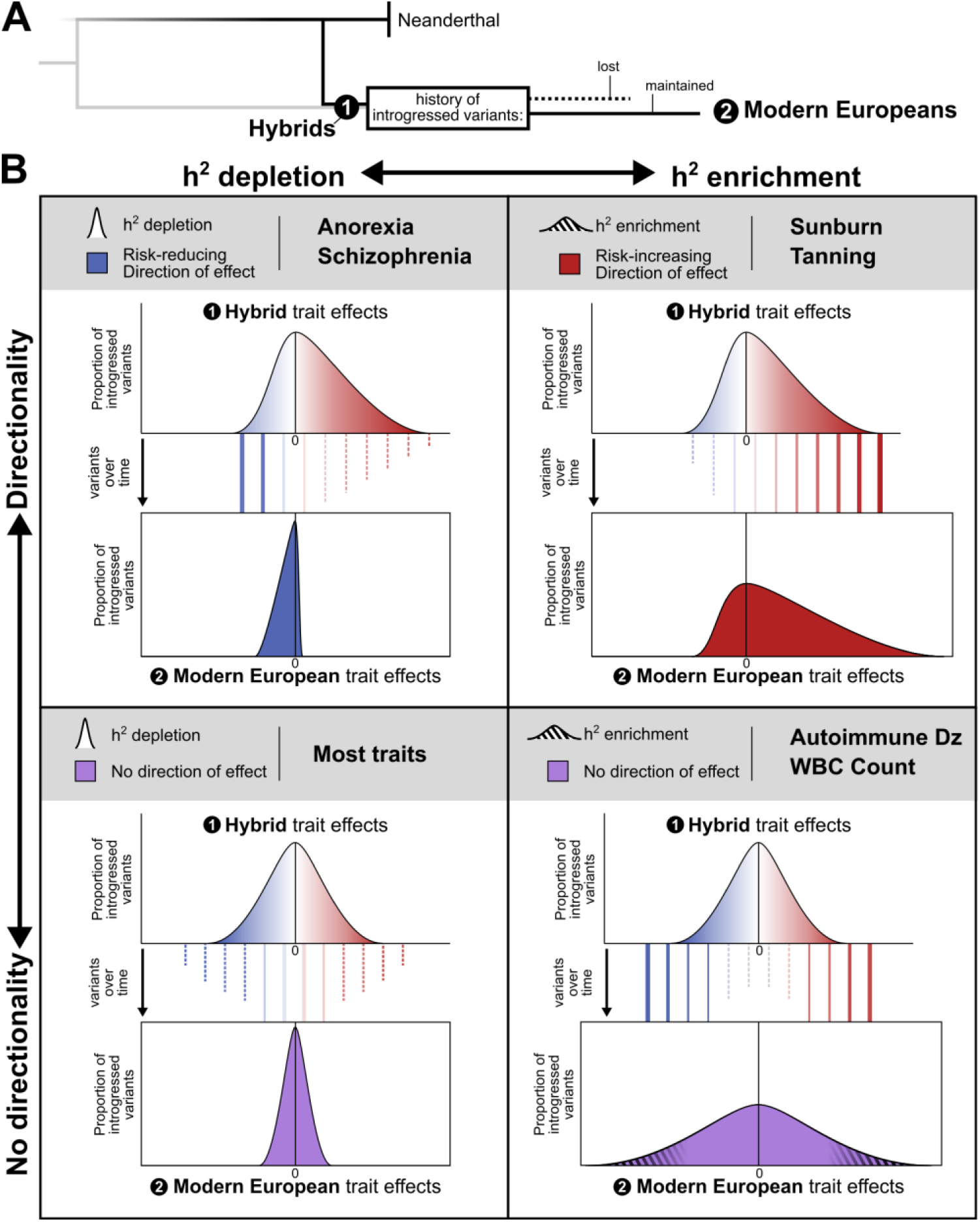
Patterns of heritability and direction of effect reveal contrasting patterns of selection on introgressed variation associated with different traits. (A) Directly after admixture, hybrid individuals had many introgressed variants (black). Selection acted on traits in hybrids as they evolved into modern Europeans causing some introgressed variants to be lost due to drift or negative selection (dashed line) and some to be maintained due to drift or positive selection (solid line). (B) In the remaining introgressed variants observed in modern Europeans, traits fall into four general quadrants on the axes of heritability enrichment vs. depletion (x-axis) and directional bias vs. no bias (y-axis). For each quadrant, we depict potential variant histories and selective pressures leading to the observed distribution of introgressed variants’ trait effects (solid and dashed lines). (Bottom Left) Heritability for most traits is depleted among introgressed variants (narrow effect distribution with most variants conferring no trait effect) and has no bias in the direction of effect (centered at zero). This suggests selection against introgressed variants that influenced these traits. (Top Right) The opposite pattern is observed in traits such as sunburn and tanning. These traits are enriched for heritability among introgressed variants (thick tail with more variants conferring trait effects than expected) and have a bias in their direction of effect (skewed). This pattern suggests that introgression introduced functional alleles that were positively selected in AMHs. (Bottom Right) Traits, like autoimmune disease risk and white blood cell (WBC) count, have heritability enrichment among introgressed variants (thick tails with many variants conferring trait effect) but no directional bias (centered). In this case, introgression likely contributed increased diversity—both trait increasing and decreasing—relevant to the trait into AMHs that was beneficial as they adapted to non-African environments. (Top Left) Finally, traits like anorexia and schizophrenia, show depletion for heritability among introgressed variants (narrow distribution), but they have a significant directionality bias in the few introgressed variants with effects (skewed). We hypothesize that this pattern could be produced by negative selection purging most introgressed alleles that influence the trait paired with selection for a small number of introgressed protective alleles.

Several limitations must be considered when interpreting our results. First, many introgressed alleles likely had pleiotropic effects and different fitness effects in modern versus archaic environments, complicating the inference of the history of selection. Second, many of the enrichments, depletions, and directionality comparisons we report are nominally significant, but do not pass strict multiple testing corrections. This is largely because the analysis methods decrease in power to partition heritability as the number of variants in the partition decreases (S-LDSC) or assess direction of effect correlations that we expect are small (signed LD profile regression). Third, recent analyses have demonstrated that estimates of enrichments are sensitive to the assumed heritability model and that variation in heritability estimates from different statistical methods are influenced by demographic factors.^72,73^ Nonetheless, our results are consistent in direction across many traits and are correlated across variant sets. Given this consistency, the small overall differences in heritability estimates in previous evaluations, and that none of our interpretations rely on magnitude of effect, we do not anticipate that using other estimation methods would dramatically change our conclusions. Fourth, we only analyze the effects of introgressed variation the context of Europeans. Further work and GWAS are need to comprehensively understand the role of introgressed variation in other (*e.g.* East and South Asian) populations. Fifth, in the direction of effect analyses, we were conservative in considering only Altai-matching alleles and expanding for LD in mapping introgressed variants to GWAS hits. Thus, some introgressed alleles with effects on traits considered may have been missed (Methods); however, our genome-wide signed LD profile regression approach considers all variants and effects. Finally, while we identify associations with between many introgressed haplotypes and traits, we do not provide molecular validation of the specific causal allele(s) behind the association.

With the growth of large cohorts including linked genotype and phenotype data, it will be valuable to extend these heritability analyses to large-scale biobank data sets from diverse populations. This will enable further quantification of the functional effects and selective pressures on introgressed variants, including introgression from Denisovans, and other alleles with unique evolutionary histories (*e.g.*, reintroduced ancestral alleles, high frequency derived alleles). We also anticipate that simulation studies can inform our understanding of the types of selective pressures required after introgression to produce the heritability patterns observed. Ultimately, increased knowledge of how remaining introgressed Neanderthal alleles influence AMH populations provides a window into understanding the phenotypic variation of Neanderthal populations over 50,000 years ago and how this variation contributed to AMH adaptation to diverse environments.

## METHODS

### Defining Neanderthal-introgressed regions

#### Larger genomic regions tolerant of Neanderthal ancestry

To select genomic regions tolerant of Neanderthal ancestry we used “segments” identified by Browning et al. 2018 using Sprime, a hidden Markov Model that identifies windows that contain a high density of SNVs in admixed populations (*i.e. Europeans*) that are not observed in unadmixed (*i.e.* African) populations.^34^ We considered the Sprime-identified segments identified using five European subpopulations (CEU, TSI, FIN, GBR, IBS). As recommended by Browning *et al.* 2018 to isolate regions with evidence of Neanderthal ancestry, we (1) considered segments identified in these five populations that have at least 30 putatively introgressed variants that could be compared to the Altai Neanderthal genome and (2) had a match rate of at least 30% to the Altai Neanderthal allele.^34^ We provide data on these sets in Fig. S1. After applying these two filters to the segments identified independently in the five European subpopulations, we merged these sets. This ultimately defines a set of segments with evidence of Neanderthal ancestry in Europeans used for the top panel of Fig. 1B. To define the second set of segments with Neanderthal ancestry minus introgressed variants (bottom panel of Fig. 1B), we identified 1000G variants in these segments and subtracted out introgressed variants (set 4 below, LD expanded to *r*^2^ > 0.5).

#### Neanderthal introgressed variants

We consider four sets of Neanderthal introgressed alleles based on Sprime analyses. From most stringent to least stringent, these sets are: (1) putatively introgressed variants identified in European subpopulations matching the Altai Neanderthal allele (used predominately in analyses in Fig. 1D-4, N = 138,774), (2) putatively introgressed variants identified in any subpopulation matching the Atlai Neanderthal allele (N = 276,902), (3) putatively introgressed variants identified in European subpopulations regardless of evidence of matching the Neanderthal allele (N = 350,577), and (4) putatively introgressed variants identified in any subpopulation regardless of evidence of matching the Neanderthal allele (used in Fig. 1C-D, N = 900,902). In sets three and four, the variants might not match the Altai Neanderthal allele at the site or a comparison might not have been possible due to lack of coverage or high confidence allele call. We present results from set one (“Altai-matching introgressed variants”) and set four (“introgressed variants”) in the main text.

#### Vernot 2016 S*-identified haplotypes and variants

For completeness, we also considered the introgressed Neanderthal haplotypes previously identified by Vernot et al. 2016.^49^ These introgressed regions were identified using the S* statistic which, like Sprime, uses mutational profiles to infer introgressed regions in the absence of any archaic reference genome. Like Sprime, S* compared mutational profiles between a putatively introgressed target populations and a non-introgressed outgroup. Sprime differs from S* in that it simultaneously considers multiple members of the target population, and Sprime allows for limited geneflow between the target population and the outgroup.

For introgressed haplotypes identified by S* in Europeans (5851), 3243 (55%) are more than 50% covered by at least one EUR segment identified by Sprime, and 2370 S* haplotypes (40%) have 0% coverage. Conversely, for introgressed segments identified by Sprime in Europeans (1733), 1128 (65%) are more than 50% covered by at least one EUR haplotype identified by S*, while 282 (16%) have 0% coverage. Therefore, Sprime segments are more of a subset of S* haplotypes than S* haplotypes are a subset of Sprime segments.

### GWAS summary statistics

#### 41 representative traits

We considered GWAS summary statistics from a previously-described representative set of 41 diseases and complex traits. ^37–48^ Previous studies using these traits had GWAS replicates (genetic correlation > 0.9) for six of these traits (BMI, Height, High Cholesterol, Type 2 Diabetes, Smoking status, Years of Education). For these six traits, we considered only the GWAS with the largest sample size so our combined analysis did not overrepresent these six. All GWAS are European-ancestry only. Many are from UK Biobank, but we note that their coding may be different than coding used in other UK Biobank heritability analyses.^37^ For example, morning person is converted into a binary variable (morning person vs. evening person) rather than the categorical ordinal scale the data was obtained on (“definitely a morning person”, “more a morning person”, “more an evening person”, “definitely an evening person”). Information on these traits are in Table S1.

#### 405 UK Biobank Traits

For a larger set of traits, we considered GWAS from the UK Biobank and 15 from FinnGen formatted for LDSC by the Neale Lab.^37,51^ For reliability of S-LDSC heritability estimates, we apply two thresholds to select GWAS based on recommendations from Finucane et al. 2015 and the Neale lab analysis.^36,51^ First, we only consider traits that meet criteria for “high” confidence estimates of SNP heritability. These traits have an effective sample size of greater than 40,000, a standard error of less than 6 times expected, and a sex bias less than 3:1. Traits with nonlinear ordinal coding of numeric values are also excluded. Second, we consider traits that are significantly heritable. These are phenotypes that have heritability estimates with *P* < 1.28×10^−12^ (z > 7). This defines a set of 405 summary statistics described in Table S2. Some traits are independent, but many of these traits are also correlated. Traits from the previous set of 41 are only included if they existed in this high-confidence set of UK Biobank/FinnGen.

#### Defining phenotypic domains

To make conclusions on a domain level, we categorize traits by their phenotypic “domains,” “chapters,” and “subchapters”. We derive these designations from the GWAS Atlas, a database of publicly available GWAS summary statistics.^53^ The GWAS Atlas had categorized many of the 405 UK Biobank traits; however, because GWAS Atlas uses different criteria for inclusion into their database, some of these traits were uncategorized. For the uncategorized UK Biobank traits and the other set of 41 traits, we manually assigned them into existing domains, chapters, subchapters based on similar categorized traits. The only edit to the existing designations was changing subchapter labels of the immunologic domain. All its subchapter instances (N=14) were labeled “Immunological System Functions.” We manually changed this generic label to either “RBC” or “WBC.” For example, reticulocyte count and mean corpuscular hemoglobin fall under RBC, while eosinophil count and neutrophil fall under WBC. The 405 GWAS cross 21 domains, 31 chapters, and 62 subchapters. However, we note that this organization is not purely hierarchical (*e.g.* some traits in the same subchapter belong in two different domains). The phenotypic domains, chapters, and subchapters assigned to each of the 405 traits are in Table S2.

### Quantifying partitioned heritability with S-LDSC

We quantified partitioned heritability using Stratified-LD Score Regression v1.0.1 (S-LDSC) to test whether an annotation of interest (*e.g.* introgressed regions or introgressed variants) is enriched for heritability of a trait.^35,36^ We use 1000 Genomes for the LD reference panel (variants with MAF > 0.05 in European samples)^74^ and HapMap Project Phase 3 (HapMap 3)^75^ excluding the MHC region for our regression variants to estimate heritability enrichment and standardized effect size metrics.

S-LDSC estimates the heritability enrichment, defined as the proportion of heritability explained by variants in the annotation divided by the proportion of all variants considered in the annotation. The enrichment of annotation *c* is estimated as

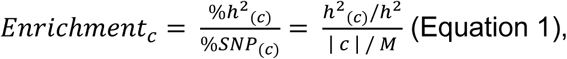

where *h*^*2*^_*(c)*_is the heritability explained by common variants in annotation *c, h*^*2*^ is the heritability explained by common variants over the whole genome, |*c*| is the number of common variants that lie in the annotation, and *M* is the number of common variants considered over the genome. Statistical significance is estimated by LDSC by using a block jacknife across adjacent variants in 200 equally-sized blocks.^36,41^

### Direction of effect: Intersection with genome-wide significant variants

To intersect introgressed variants with genome-wide significant variants, we first used PLINK to LD expand the Altai-matching introgressed Neanderthal variants (set 1, described in “Neanderthal introgressed variants” methods section) to perfect LD (*r*^2^ > 0.999).^76^ LD was calculated for variants within 1 Mb of each introgressed variant using the 1000G European reference population while preserving the “phase” of the allele in LD with the Neanderthal allele.^74^ We eliminated any duplicates (*i.e.* if two introgressed variants in perfect LD were both tagging another variant). We intersected this LD-expanded set of introgressed variants with the GWAS summary statistics using rsIDs. We oriented the sign of the summary statistic (the z-score) relative to the archaic allele (or the allele in perfect LD to the archaic allele). For example, if a variant is positively associated with a trait (z-score is +6 with GWAS effect allele “A” and alternative allele “C”), but the archaic allele is “C”, we flip the z-score to be -6 because the archaic allele “C” is negatively associated with the trait.

For eight traits (AutoimmuneDz, Balding, Sunburn, FVC, Heel_T_Score, MorningPerson, MenopauseAge, WBCCount), we filtered the introgressed variant-summary statistic intersection at different thresholds of genome-wide significance (*P* < 1 ×10^−8^, *P* < 5 ×10^−8^, *P* < 1 ×10^−6^). We then pruned variants at various levels of LD (*r*^2^ = 1, *r*^2^ = 0.8, *r*^2^ = 0.5) to reduce redundant counts due to linked loci. We used the LDmatrix tool in the LDlink API to calculate the pairwise LD to prune linked variants (with the 1000G EUR as a reference).^77^ We then counted the number of introgressed alleles associated with the positive and the negative directions of the trait. With quantified significance with a chi-squared goodness of fit test.

#### Limitations of genome-wide significant variant intersection

We caution overinterpretation of these results and highlight some of the limitations of this method. First, despite LD expansion, only 29% of the introgressed alleles could be intersected with variation interrogated by the GWAS (LD expanded to *r*^2^ = 1). Therefore, this analysis would not be able to investigate directionality of introgressed variants in regions not perfectly tagged by the genotyping array used for the GWAS. However, 61% of the Sprime segments (larger windows tolerant of Neanderthal introgression) have at least one introgressed variant interrogated by the GWAS; therefore, we feel confident that this analysis is sampling broadly across introgressed regions. Second, by considering only genome-wide significant variants, this directionality analysis limits to loci in the extremes of the GWAS distribution. It does not consider the global genome-wide relationship between introgressed alleles and directionality of trait-associated variation at varying levels of effect size and significance. However, we show these results are consistent at a less stringent level of genome-wide significance (*P* < 5 ×10^−8^, Fig. S7).

### Direction of effect: Signed LD profile regression (SLDP) analysis

SLDP quantifies the genome-wide directional effect of a signed functional annotation on polygenic disease risk. To do so, SLDP calculates a correlation between a vector of variant effects on a trait (from GWAS summary statistics, α^) and a vector of those variants’ aggregate tagging of an annotation (*R*ν).^55^ Our annotation is each variant’s maximum LD to a Neanderthal introgressed allele (we term *Neanderthal LD profile*). This allows us to quantify if there is a genome-wide relationship between a variant’s LD to a Neanderthal allele and the direction of that variant’s trait association. This is distinct from previous stratified-LD score regression (S-LDSC) analyses because S-LDSC quantifies heritability enrichment in an annotation of interest independent of directionality.

More specifically, SLDP regresses α^ (the vector of marginal correlations between variant alleles and a trait) on vector ν (our signed functional annotation) to estimate *r*_*f*_, the functional correlation between the annotation and trait, using

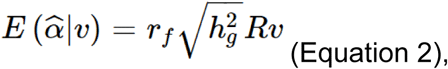

where *R* is the LD matrix from the 1000G Phase 3 European reference, *h*^*2*^ is the trait’s SNP-heritability. Together, *R*ν is a vector quantifying each variant’s aggregate tagging of the annotation, termed the *signed LD profile.* SLDP uses generalized least-squares regression across HapMap 3 variants excluding the MHC region (M = 1,187,349). It also conditions the regression on a “signed background model” that quantifies the directional effects of minor alleles to reduce confounding due to genome-wide negative selection or population stratification (using five equally sized MAF binds). False discovery rates and P-values are obtained by empirically generating a null disruption through randomly flipping the signs of ν in large blocks. For a detailed description of the SLDP method, derivation, estimands, and validation, see Reshef *et al.* 2018.^55^

We conduct this SLDP analysis on the same eight GWAS summary statistics considered in the previous direction of effect analysis (AutoimmuneDz, Balding, Sunburn, FVC, Heel_T_Score, MorningPerson, MenopauseAge, WBCCount). To generate our functional annotation, we used PLINK to calculate pair-wise LD between the Altai-matching introgressed variants (set 1, described in “Neanderthal introgressed variants” methods section) and the 1000 Genomes Phase 3 European reference panel (∼10M variants).^74^ We considered LD limited to variant pairs within 1 Mb and *r*^2^ > 0.2. For each variant in the reference panel, the annotation (ν) is the maximum *r*^2^ value to the Neanderthal variants. The input annotation (ν) is generated in reference to the allele that is on the Neanderthal haplotype. However, for the SLDP regression, the signs (for both α^ and Rv) are oriented in reference to the European minor allele.

For interpretability of the visualizations, all plots show α^ and *R*ν with reference to the Neanderthal allele. For example, if Neanderthal variant X is LD (and in-phase) with SLDP regression variant Y at *r*^2^ = 0.5, variant Y’s functional annotation (ν) is 0.5. We plot the sign of α^ (from the GWAS) with reference to Y as the effect allele (A1). All plots describing SLDP results display the residuals of α^ (y-axis) and *R*ν (x-axis) for each variant. This residual reflects that all analyses are conditioned on the “signed background model” described above.

### Identifying genomic windows with association Neanderthal LD profile and trait effect

To locate regions of interest, we identify genomic windows with a strong correlation between LD to introgressed alleles and trait-associated risk or protection. From the per-variant output from SLDP regression (M = 1,187,349), we calculate Pearson correlation coefficients between the residuals of α^ and *R*ν for 30 kb sliding windows centered around each SLDP regression variant. We select windows that have at least 15 SLDP regression variants and an *r*^2^ > 0.5 (correlation in either direction). We join overlapping windows; therefore, final windows are often bigger than 30 kb and can have a correlation coefficient less than 0.5. We only consider windows that have at least one variant marginally associated with the trait (*P <* 1×10^−4^) and windows that overlap at least one Altai-matching Neanderthal introgressed allele (set 1; see above). Table S6 gives these windows for the eight traits considered by SLDP analyses.

Figures depicting the windows of interest identified were generated using the UCSC Genome Browser.^78^ eQTL and sQTL analysis and plots were generated using the Genotype-Tissue Expression (GTEx) Project (V8 release) Portal on 4/29/2020.^59^ GTEx V8 results are in GRCh38 and were lifted over to GRCh37 (hg19) for comparison with the windows of interest. PheWAS results are from the GWAS Atlas and consider 4756 traits.^53^

### Data analysis and figure generation

All genomic coordinates and analysis refer to *Homo sapiens* (human) genome assembly GRCh37 (hg19) unless otherwise specified. Data and statistical analyses were conducted using Python 3.5.4 (Anaconda distribution), R 3.6.1, Jupyter Notebook, BedTools v2.26, and PLINK 1.9.^76,79^ Figure generation was significantly aided by Matplotlib, Seaborn, and Inkscape.^80–82^.

### DATA AVAILABILITY

The publicly available datasets and software used for analysis are available in the following repositories: introgressed variants and segments from Sprime [https://data.mendeley.com/datasets/y7hyt83vxr]^34^, introgressed variants and segments from S* [http://akeylab.gs.washington.edu/vernot_et_al_2016_release_data/introgressed_tag_snp_frequencies/]^49^, LDSC [https://github.com/bulik/ldsc]^35,36^, GWAS traits formatted for LDSC from the Alkes Price lab [https://data.broadinstitute.org/alkesgroup/LDSCORE/independent_sumstats/], UK Biobank traits formatted for LDSC from the Benjamin Neale lab [http://www.nealelab.is/uk-biobank]^51^, GWAS Atlas [https://atlas.ctglab.nl/]^53^, Signed LD Profile Regression [https://github.com/yakirr/sldp]^55^, and the GTEx Project Portal [https://gtexportal.org/home/]^59^.

The datasets we generated are available in the trait-h2-neanderthals GitHub repository [https://github.com/emcarthur/trait-h2-neanderthals]. They include bed files of all genomic partitions considered (regions with Neanderthal ancestry, sets of introgressed variants), all results of partitioned heritability analysis output (for the 41 traits formatted from the Price Lab and the 405 traits from the UKBioBank formatted by the Neale Lab) and signed LD profile regression results. The repository also contains a Jupyter notebook with python code used for data analysis and all figure generation.

## Authors’ contributions

EM, DR, and JAC conceived and designed all the work presented here. EM and DR conducted all the analysis. EM and JAC interpreted the results, drafted the work, and substantively revised the manuscript. EM, DR, and JAC have approved the submitted version and have agreed to be accountable for their contributions.

## Acknowledgements

The authors would like to thank Abin Abraham, Laura L. Colbran, Sarah Fong, Lea K. Davis and other members of the Capra Lab for helpful discussions and manuscript comments. This work was conducted in part using the resources of the Advanced Computing Center for Research and Education (ACCRE) at Vanderbilt University, Nashville, TN.

