## Supplementary material for "Quantifying the contribution of Neanderthal introgression to the heritability of complex traits": Figures S1-S8, Tables S1,S3-S5

### SUPPLEMENTAL FIGURES

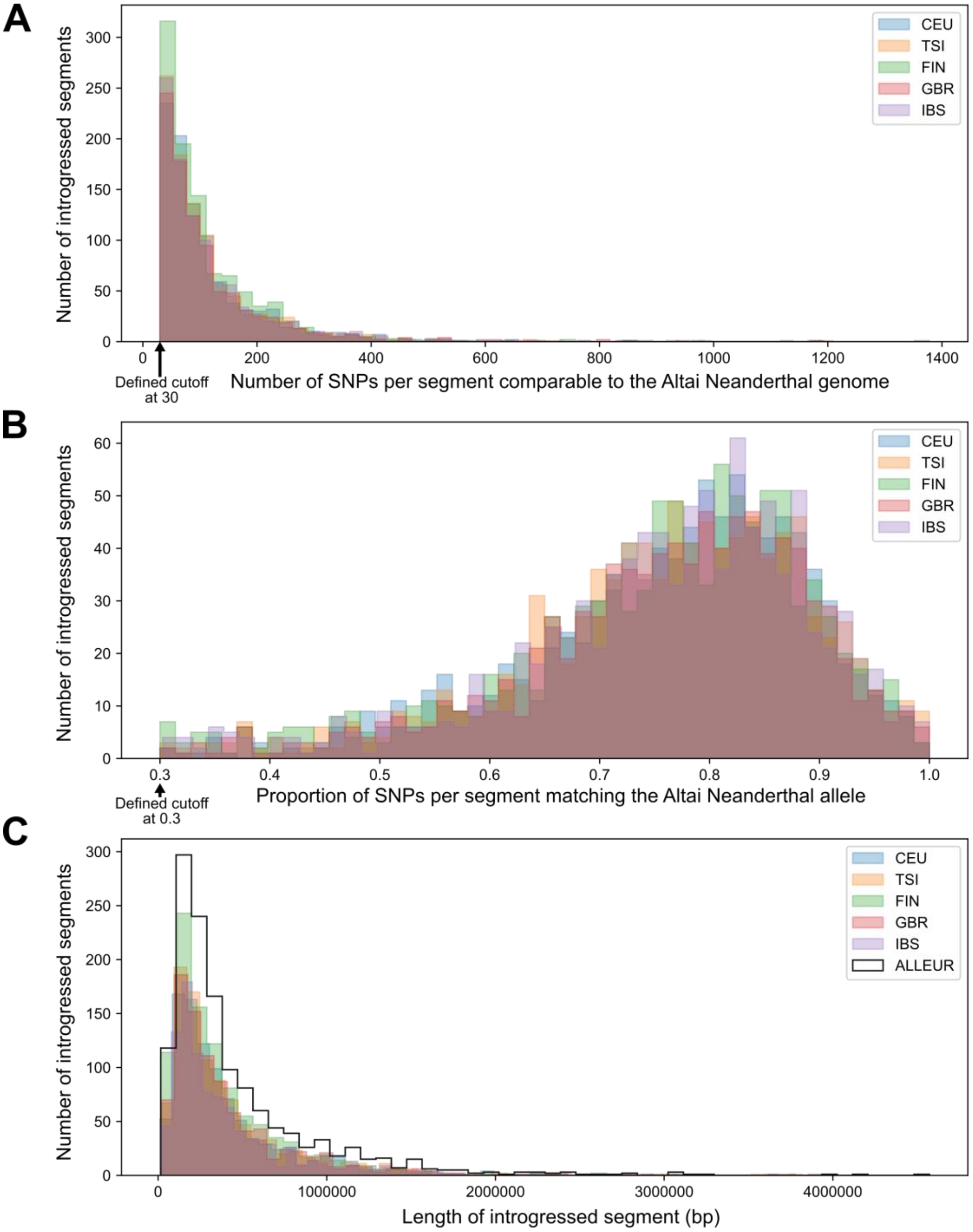


**Figure S1. Defining genomic regions tolerant of Neanderthal ancestry.** Of the introgressed segments defined by Browning et al. 2018 we consider those observed in any of the European subpopulations (CEU, TSI, FIN, GBR, IBS), (A) that have at least 30 putative introgressed variants that are comparable to the Altai Neanderthal genome (after filtering, these segments have an average of 116 comparable variants) and (B) that these putative introgressed variants have at least 30% match to the Altai Neanderthal allele. After filtering, these segments have a 76% match on average. (C) The size distribution of these independently identified segments after applying these two filters. We also consider the union of these sets (black). Ultimately, we define 1345 segments that have a median length of 299 kb (IQR: 174 – 574 kb). This set is used in Fig. 1B. (CEU: Utah Residents with Northern and Western European Ancestry, TSI: Toscani in Italia, FIN: Finnish in Finland, GBR: British in England and Scotland, IBS: Iberian Population in Spain)

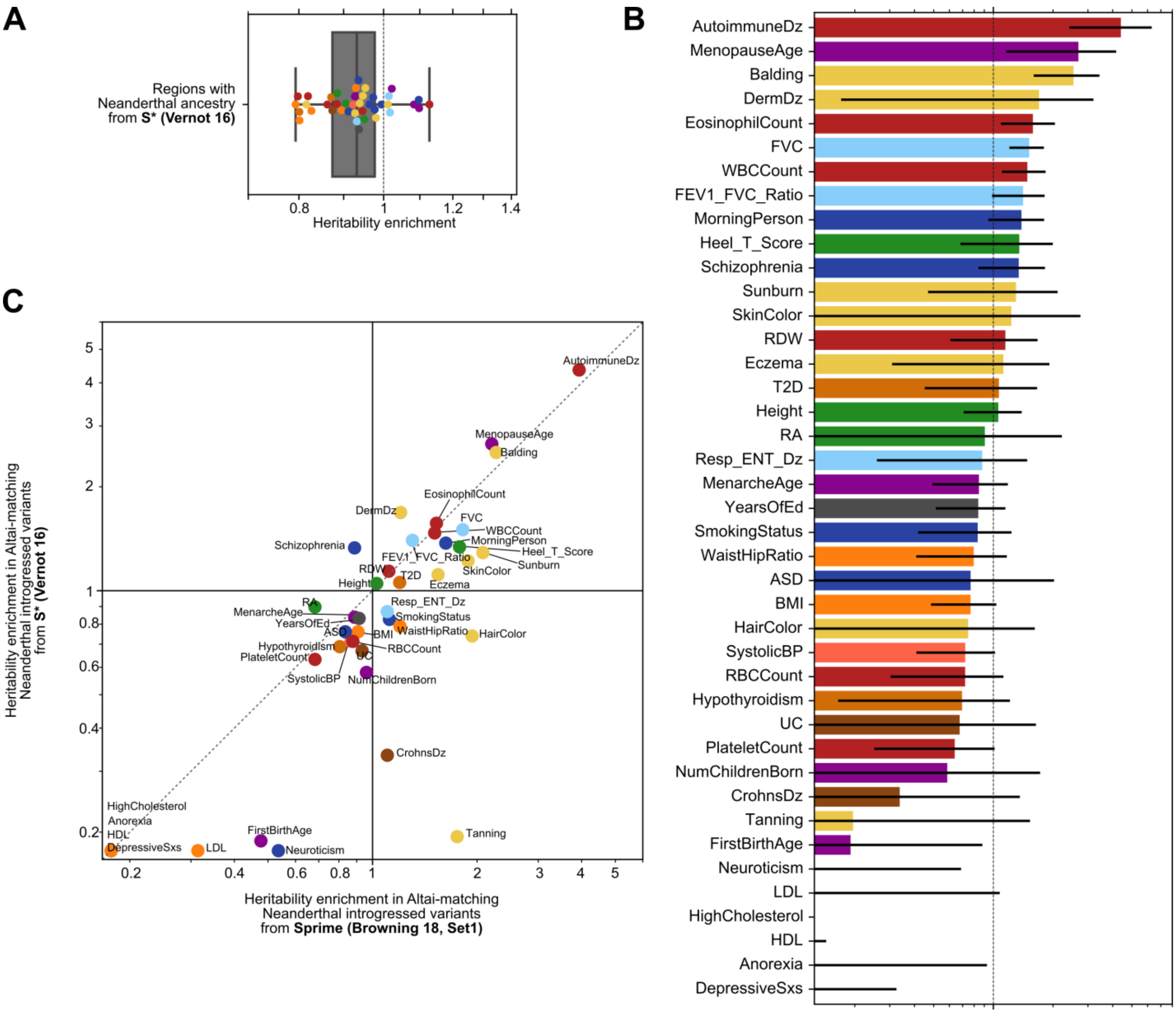


**Figure S2. Trait heritability patterns in regions tolerant of Neanderthal ancestry and introgressed variants are consistent when defined based on variants identified by S* from Vernot *et al.* 2016.** (A) Similar to Fig 1B, we show that traits are broadly depleted of heritability in regions tolerant of Neanderthal ancestry defined using haplotypes from S* (0.93x background expectation, $P = 1\times10^{-5}$). (B) For a set of Altai-matching Neanderthal introgressed variants identified by S*, we show the trait-by-trait partitioned heritability analysis. Error bars represent standard errors and traits heritability depletion less than 0.125 are truncated. This set includes 132,296 variants and is comparable to the Altai-matching “set 1” variants identified by Sprime (N = 138,774) which is shown in Fig, S2A. (C) Trait heritability compared between Sprime-identified Altai-matching “set 1” introgressed variants (x-axis) and S*-identified high-confidence variants (y-axis), are highly correlated ($r^2 = 0.79$).

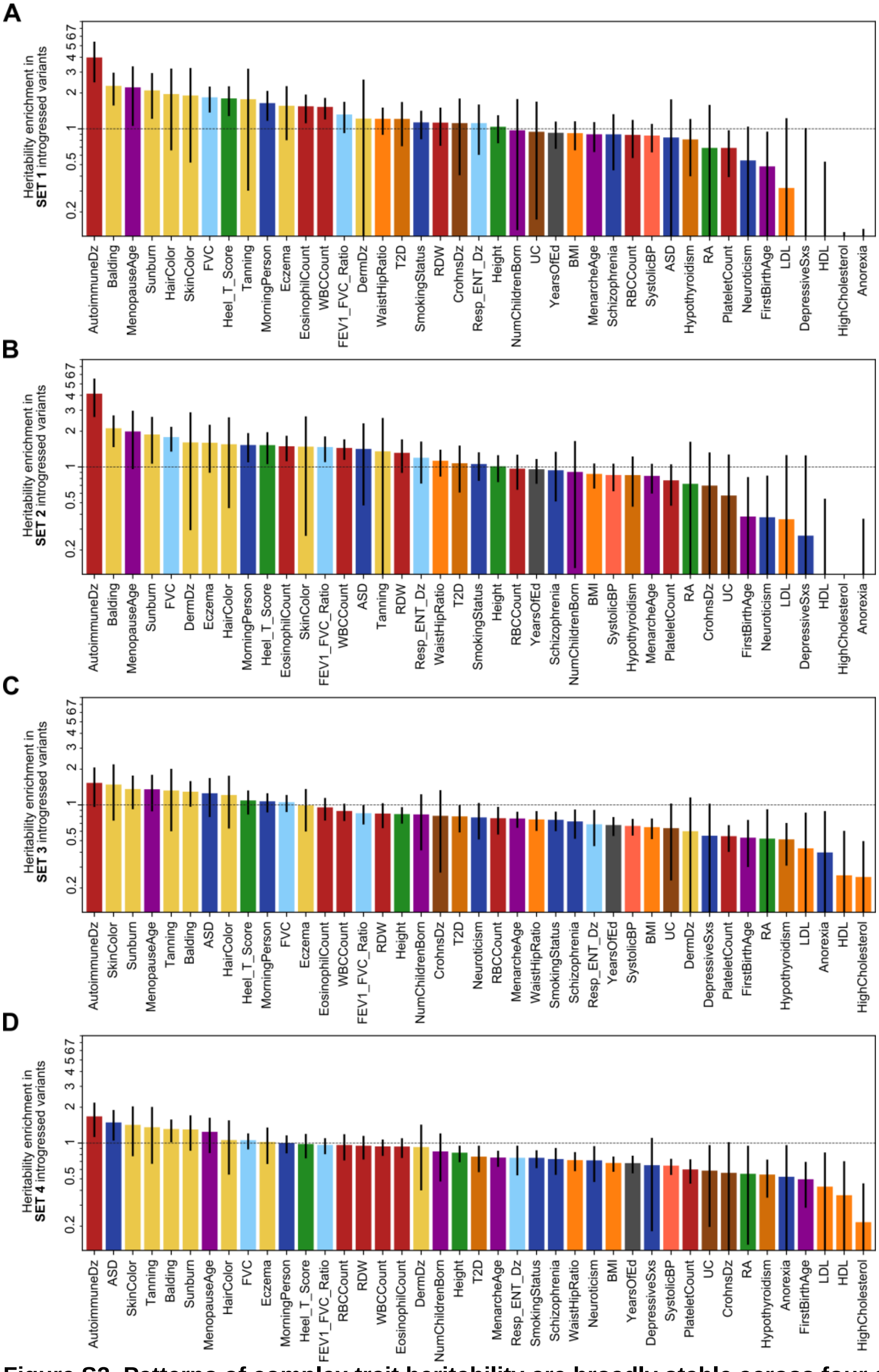


**Figure S3. Patterns of complex trait heritability are broadly stable across four different sets of Neanderthal introgressed variation.** From the most stringent set of Altai-matching variants observed in Europeans (set 1, A) to the most inclusive set of introgressed variants observed in any subpopulation (set 4, D), we show the heritability enrichment (or depletion) ordered by magnitude. Set 4 (D) is the same as Fig. 1C. The relationship between Set 4 (D) and Set 1 (A) is shown in Fig. 1D. Details of each set are in the methods. Error bars represent standard errors. Traits with depletion less than 0.125 are truncated.

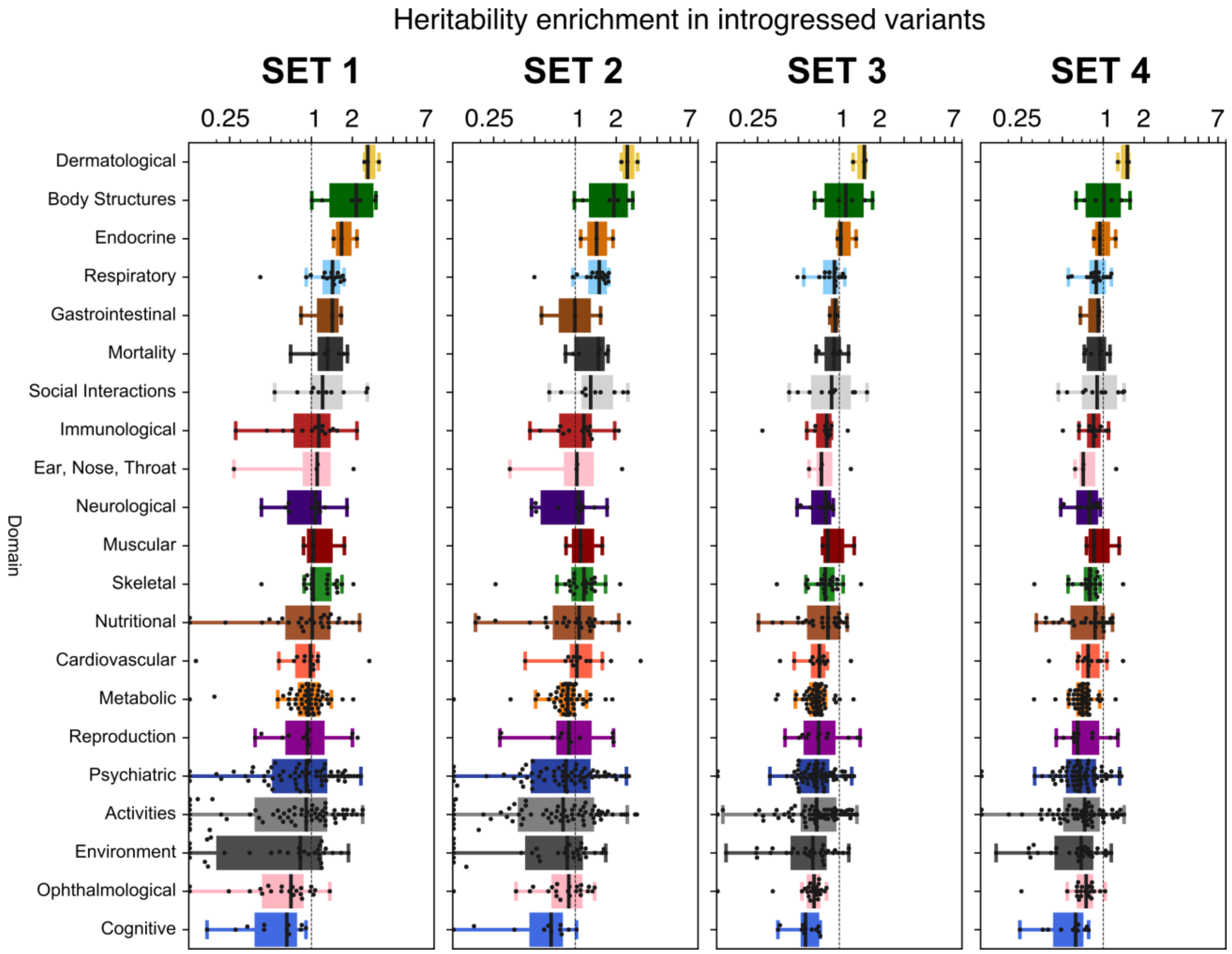


**Figure S4. Patterns of complex trait heritability across 405 traits organized by DOMAIN across four different sets of Neanderthal introgressed variation.** Across four sets of Neanderthal introgressed variation (from most stringent to least stringent [Methods]), we show the trait heritability enrichment (or depletion) across 21 phenotypic domains. Domains are ordered by magnitude of the median enrichment in Set 1 variants for comparison across sets. Results from Set 1 are the same as those depicted in Fig. 2A. Each point represents heritability enrichment or depletion of one trait in Altai-matching introgressed variants. Traits with depletion less than 0.125 are plotted on the baseline for visualization.

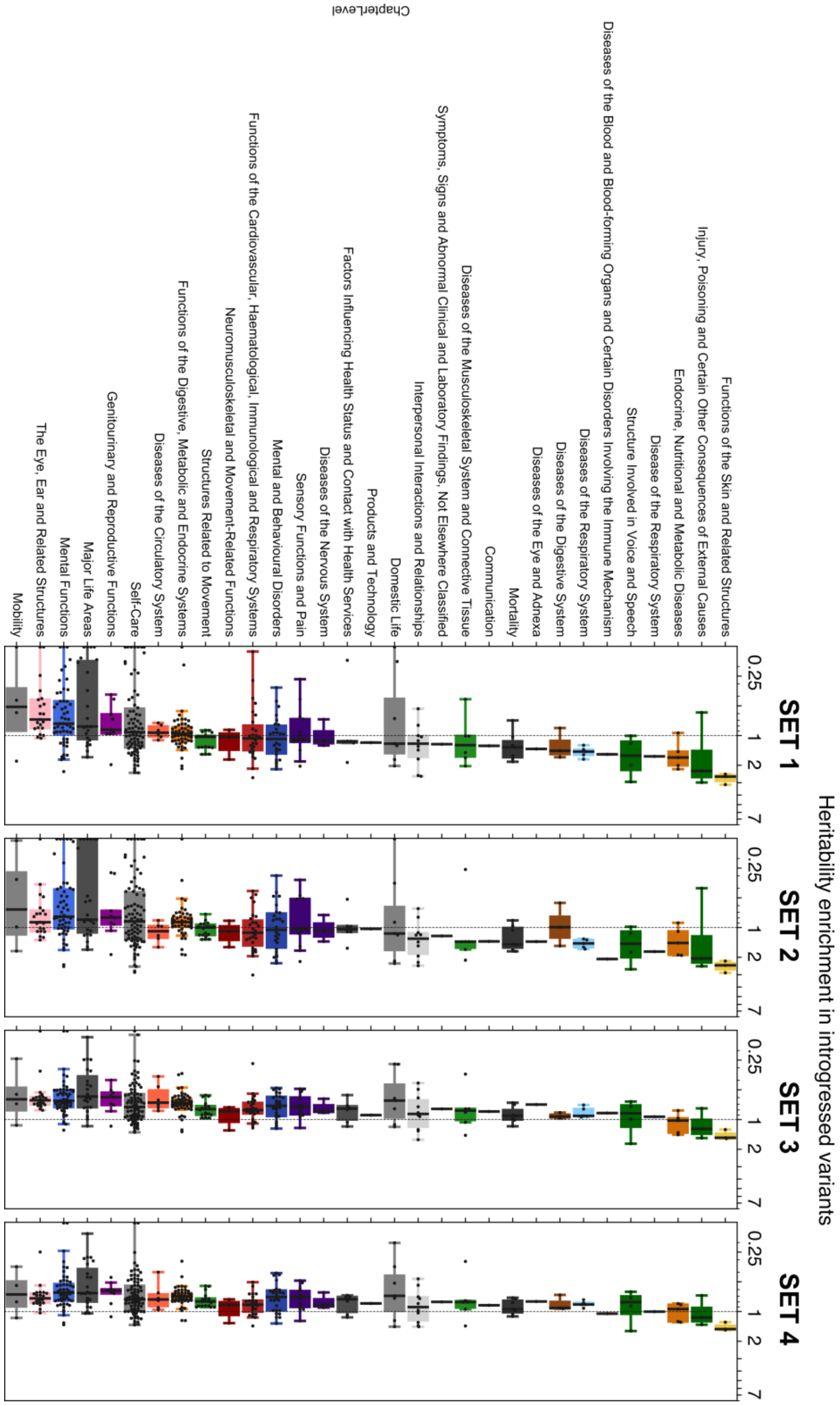


**Figure S5. Patterns of complex trait heritability across 405 traits organized by CHAPTER across four different sets of Neanderthal introgressed variation.** Across four sets of Neanderthal introgressed variation (from most stringent to least stringent [Methods]), we show the trait heritability enrichment (or depletion) across 31 phenotypic chapters. Chapters are ordered by magnitude of the median enrichment in Set 1 variants for comparison across sets. Each point represents heritability enrichment or depletion of one trait in Altai-matching introgressed variants. Traits with depletion less than 0.125 are plotted on the baseline for visualization.

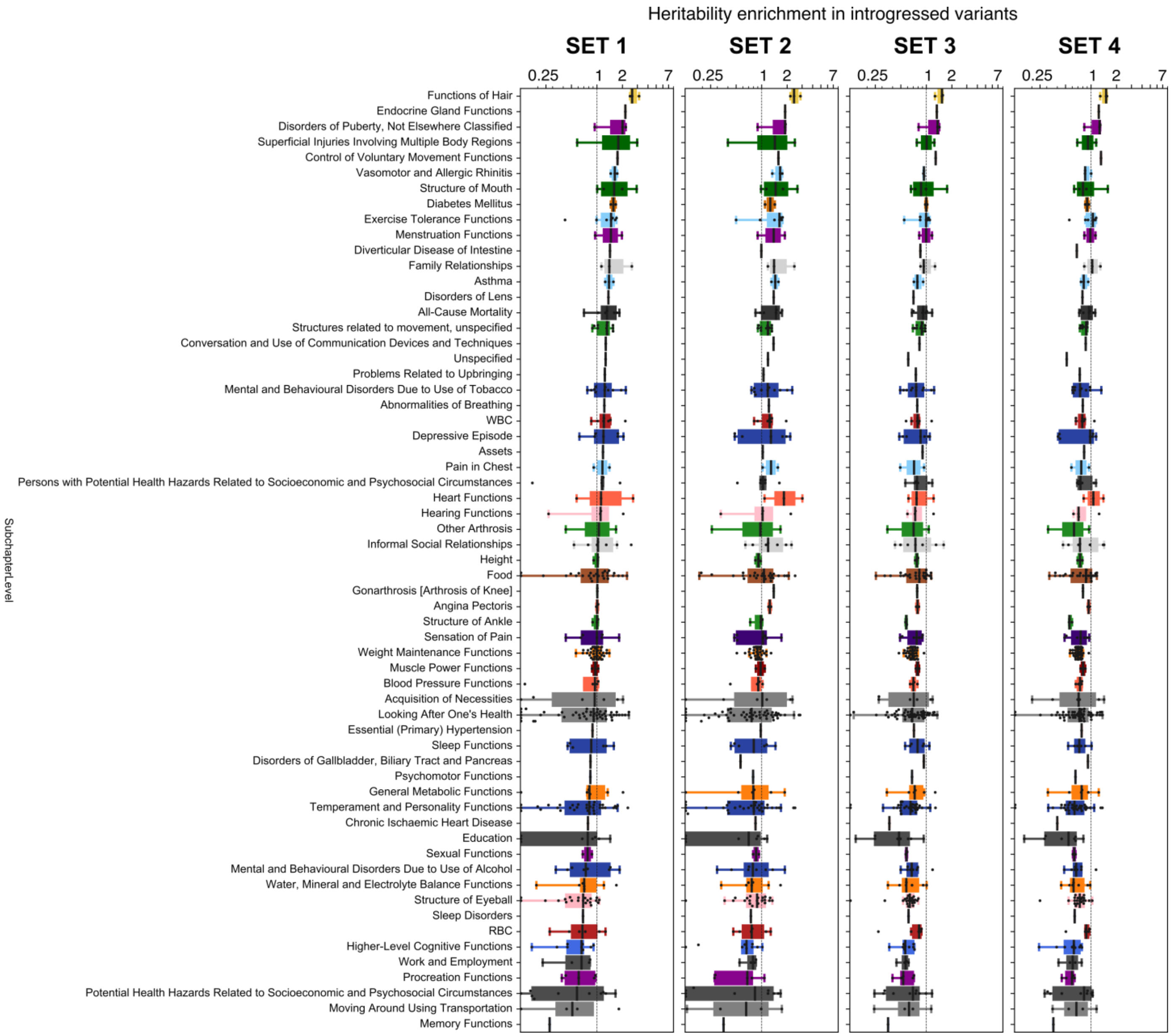


**Figure S6. Patterns of complex trait heritability across 405 traits organized by SUBCHAPTER across four different sets of Neanderthal introgressed variation.** Across four sets of Neanderthal introgressed variation (from most stringent to least stringent [Methods]), we show the trait heritability enrichment (or depletion) across 62 phenotypic subchapters. Subchapters are ordered by magnitude of the median enrichment in Set 1 variants for comparison across sets. A subset of the results from Set 1 are the same as those depicted in Fig. 2B-E. Each point represents heritability enrichment or depletion of one trait in Altai-matching introgressed variants. Traits with depletion less than 0.125 are plotted on the baseline for visualization.

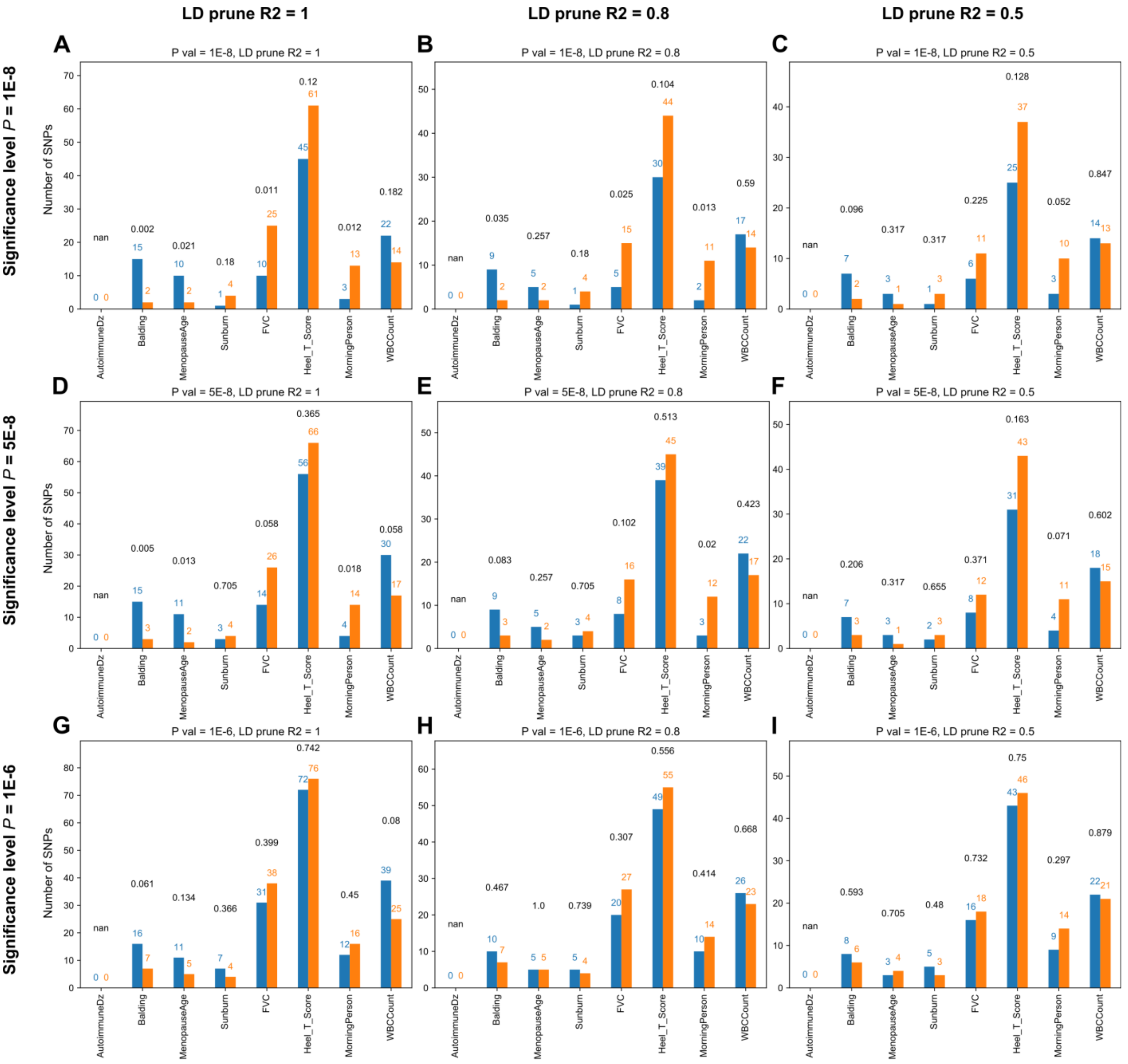


| | AutoimmuneDz | Balding | MenopauseAge | Sunburn | FVC | Heel_T_Score | MorningPerson | WBCCount |
|---|---|---|---|---|---|---|---|---|
| Direction of effect | protection | hair loss | younger | never burnt | smaller | less dense | evening | lower |
| | risk | full hair | older | sunburns | larger | more dense | morning | higher |

**Figure S7. For introgressed variants with the strongest trait-associated effects, directionality is stable even at different significance levels and pruning thresholds.** For eight traits, we intersected introgressed Altai-matching Neanderthal alleles (LD-expanded to $r^2 = 1$) with the genome-wide significant variants from each GWAS. After pruning for linked variants, we plot the number of significantly associated introgressed variants by their direction of effect (risk-increasing or risk-decreasing [legend]). Fig. 3A shows this result for the genome-wide significant threshold $P < 1 \times 10^{-8}$ and pruning threshold of $r^2 = 1$. Here, we show these results are consistent at different genome-wide significance thresholds ([A, B, C] $P < 1 \times 10^{-8}$; [D, E, F]; $P < 5 \times 10^{-8}$; [G, H, I] $P < 1 \times 10^{-6}$) and different LD pruning thresholds ([A, D, E] $r^2 = 1$; [B, E, H] $r^2 > 0.8$; [C,F,I] $r^2 > 0.5$). Black numbers above the bars represent $P$ values ($\chi^2$ goodness of fit test).

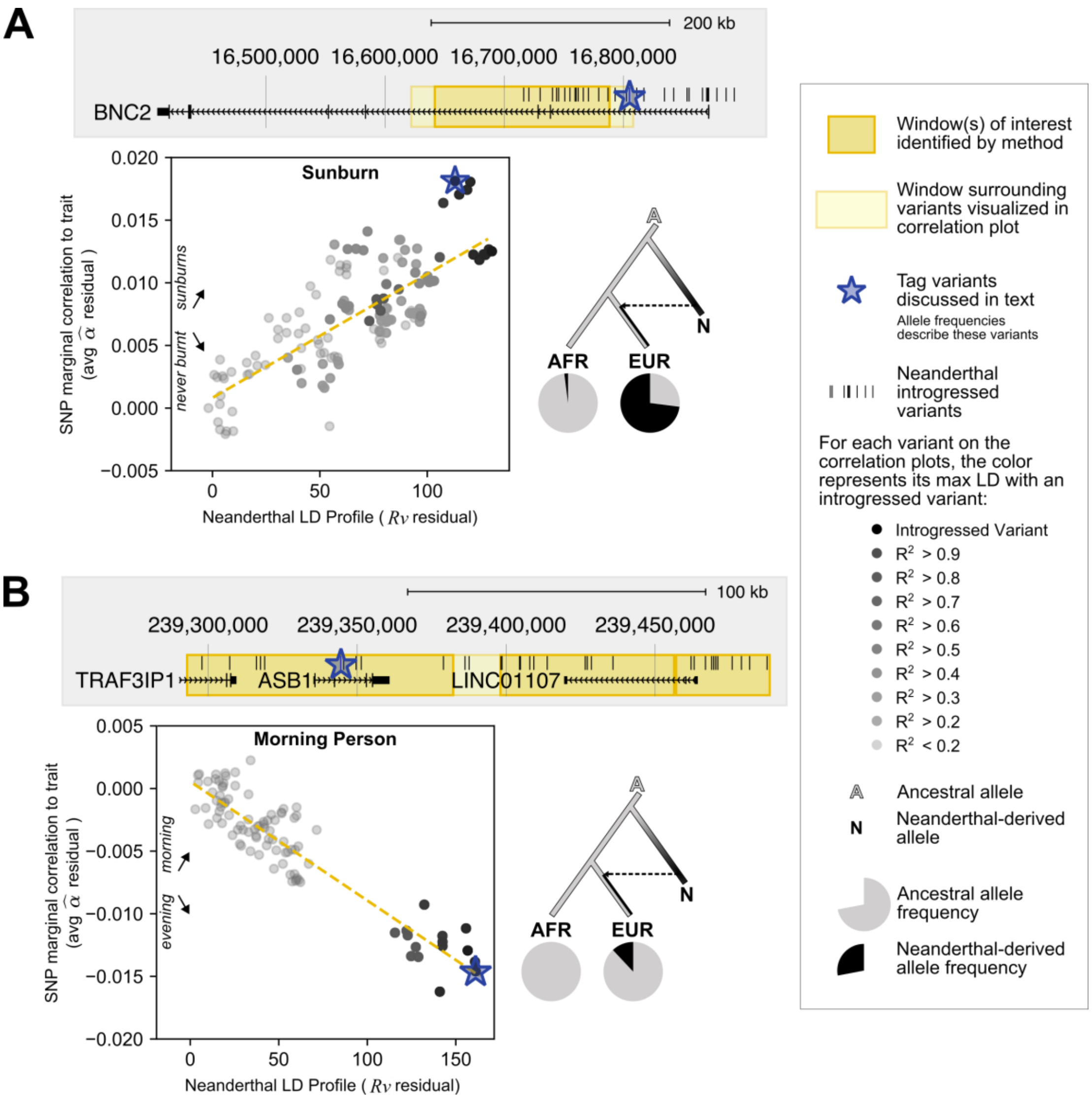


**Figure S8. Windows with strong correlations between Neanderthal LD profile and trait-association highlights known genes implicated in introgression's effect on sunburn risk and chronotype.** (A) The genomic window (chr9:16,641,651-16,787,775) overlaps *BNC2* and has a positive relationship (R = +0.82) between Neanderthal LD profile and sunburn risk. Supporting this association, the starred variant (rs10962612; EUR AF: 0.73; AFR AF: 0.02) was previously shown to tag an introgressed haplotype and associate with childhood sunburn risk and poor tanning.[4] (B) The genomic region shown highlights three windows (chr2:239,292,973-239,382,296, chr2:239,398,170-239,456,308, chr2:239,457,097-239,488,435) around *ASB1* with a negative relationship between Neanderthal LD profile and morning person status. The scatter plot shows this negative correlation (R = -0.92) for chr2:239,292,973-239,488,435; hence, increased Neanderthal LD profile is associated with increased eveningness. Supporting this association, the starred variant (rs3191996; EUR AF:0.12; AFR AF; 0.00) was previously identified as an archaic allele associated with preference for being an evening person.[4] For each example, we display the genomic region overlapped by the identified window(s) of interest (dark yellow box), genes, all Altai-matching Neanderthal-introgressed variants (black marks). For the region in light yellow, we display a scatter plot between the variant's Neanderthal LD Profile (*R*v) and trait marginal correlation (α̂). Each variant is colored by its maximum LD to an introgressed variant. We display the evolutionary history (dendrogram) of each discussed tag variant (blue star) with its allele frequency in Africans (AFR) and Europeans (EUR)(pie charts).

### SUPPLEMENTAL TABLES

**Table S1.** Genome-wide association study (GWAS) traits used for partitioned heritability analyses with S-LDSC

| Nickname | Trait | M | h2 | h2_SE | N | Source |
|---|---|---|---|---|---|---|
| Anorexia | Anorexia | 931184 | 0.2153 | 0.0169 | 32143 | Boraska et al. 2014 Mol Psych[42] |
| ASD | Autism_Spectrum | 1173307 | 0.4607 | 0.0517 | 10263 | PGC Cross-Disorder Group, 2013 Lancet[43] |
| AutoimmuneDz | Auto_Immune_Traits_(Sure) | 1187056 | 0.0068 | 0.0013 | 459324 | UKBiobank[37] |
| Balding | Balding_Type_I | 1187056 | 0.2154 | 0.019 | 208336 | UKBiobank[37] |
| BMI | BMI | 1187056 | 0.252 | 0.0071 | 457824 | UKBiobank[37] |
| CrohnsDz | Crohn's_Disease | 1051514 | 0.4723 | 0.0575 | 20883 | Jostins et al., 2012 Nature[44] |
| DepressiveSxs | Depressive_symptoms | 1115393 | 0.0473 | 0.0037 | 161460 | Okbay et al., 2016 Nat Genet[45] |
| DermDz | Dermatologic_Diseases | 1187056 | 0.0094 | 0.0014 | 459324 | UKBiobank[37] |
| Eczema | Eczema | 1187056 | 0.0675 | 0.0038 | 458699 | UKBiobank[37] |
| EosinophilCount | Eosinophil_Count | 1187056 | 0.1977 | 0.0143 | 439938 | UKBiobank[37] |
| FEV1_FVC_Ratio | FEV1-FVC_Ratio | 1187056 | 0.2336 | 0.0113 | 371949 | UKBiobank[37] |
| FirstBirthAge | Age_first_birth | 1079424 | 0.0617 | 0.0033 | 222037 | Barban et al., 2016 Nat Genet[46] |
| FVC | Forced_Vital_Capacity_(FVC) | 1187056 | 0.2068 | 0.0065 | 371949 | UKBiobank[37] |
| HairColor | Hair_Color | 1187056 | 0.4523 | 0.1497 | 452720 | UKBiobank[37] |
| HDL | HDL | 1019272 | 0.1362 | 0.0166 | 99900 | Teslovich et al., 2010 Nature[47] |
| Heel_T_Score | Heel_T_Score | 1187056 | 0.3628 | 0.0307 | 445921 | UKBiobank[37] |
| Height | Height | 1187056 | 0.6034 | 0.027 | 458303 | UKBiobank[37] |
| HighCholesterol | High_Cholesterol | 1187056 | 0.0468 | 0.0039 | 459324 | UKBiobank[37] |
| Hypothyroidism | Hypothyroidism | 1187056 | 0.0459 | 0.0037 | 459324 | UKBiobank[37] |
| LDL | LDL | 1017973 | 0.121 | 0.0166 | 95454 | Teslovich et al., 2010 Nature[47] |
| MenarcheAge | Age_at_Menarche | 1187056 | 0.2457 | 0.0102 | 242278 | UKBiobank[37] |
| MenopauseAge | Age_at_Menopause | 1187056 | 0.1215 | 0.0086 | 143025 | UKBiobank[37] |
| MorningPerson | Morning_Person | 1187056 | 0.1002 | 0.0035 | 410520 | UKBiobank[37] |
| Neuroticism | Neuroticism | 1187056 | 0.1113 | 0.0037 | 372066 | UKBiobank[37] |
| NumChildrenBorn | Number_children_ever_born | 1080059 | 0.0256 | 0.0018 | 318863 | Barban et al., 2016 Nat Genet[46] |
| PlateletCount | Platelet_Count | 1187056 | 0.349 | 0.0294 | 444382 | UKBiobank[37] |
| RA | Rheumatoid_Arthritis | 1125155 | 0.1694 | 0.023 | 38242 | Okada et al., 2014 Nature[48] |
| RBCCount | Red_Blood_Cell_Count | 1187056 | 0.2434 | 0.0191 | 445174 | UKBiobank[37] |
| RDW | Red_Blood_Cell_Distribution_Width | 1187056 | 0.2234 | 0.0198 | 442700 | UKBiobank[37] |
| Resp_ENT_Dz | Respiratory_and_Ear-nose-throat_Diseases | 1187056 | 0.0483 | 0.0034 | 459324 | UKBiobank[37] |
| Schizophrenia | Schizophrenia | 1083014 | 0.4512 | 0.0189 | 70100 | SCZ Working Group of the PGC, 2014 Nature[39] |
| SkinColor | Skin_Color | 1187056 | 0.1896 | 0.0539 | 453609 | UKBiobank[37] |
| SmokingStatus | Smoking_Status | 1187056 | 0.0972 | 0.0032 | 457683 | UKBiobank[37] |
| Sunburn | Sunburn_Occasion | 1187056 | 0.0915 | 0.0162 | 344229 | UKBiobank[37] |
| SystolicBP | Systolic_Blood_Pressure | 1187056 | 0.1966 | 0.007 | 422771 | UKBiobank[37] |
| T2D | Type_2_Diabetes | 1187056 | 0.043 | 0.0025 | 459324 | UKBiobank[37] |
| Tanning | Tanning | 1187056 | 0.172 | 0.0609 | 449984 | UKBiobank[37] |
| UC | Ulcerative_Colitis | 1076834 | 0.2424 | 0.032 | 27432 | Jostins et al., 2012 Nature[44] |
| WaistHipRatio | Waist-hip_Ratio | 1187056 | 0.1423 | 0.0067 | 458417 | UKBiobank[37] |
| WBCCount | White_Blood_Cell_Count | 1187056 | 0.1873 | 0.0105 | 444502 | UKBiobank[37] |
| YearsOfEd | College_Education | 1187056 | 0.1299 | 0.0037 | 454813 | UKBiobank[37] |

**Table S2.** See tab in SupplementalTables.xlsx
This table contains details about each of the 405 trait GWASs used in the analysis for Figure 2. The GWASs are from UK Biobank and FinnGen. The formatting for LDSC, heritability results for each phenotype, and GWAS metadata were organized by the Neale Lab website.[37,51,52] The Domain, Chapter, and Subchapter labels are from the GWAS Altas.[53]

**Table S3.** For each of the phenotypic domains, we list the median and mean heritability enrichment. These results are also plotted in Fig. 2A. For those domains which are depleted (enrichment below 1), we also report the fold-depletion (1/Enrichment). Domains are ordered by their median enrichment. The mean enrichment was calculated on the log-transformed enrichment values. For enrichments that were calculated to be less than zero by S-LDSC (because of high variability in estimates in regions with low heritability), we assigned them the lowest positive enrichment value (0.000497) before log-transforming to calculate the mean enrichment.

| Domain | Enrichment (median) | Fold-depletion (median) | Enrichment (mean) | Fold-depletion (mean) | P | N (traits) |
|---|---|---|---|---|---|---|
| **Dermatological** | 2.602 | NA | 2.719 | NA | 0.006 | 3 |
| **Body Structures** | 2.135 | NA | 1.907 | NA | 0.018 | 6 |
| **Endocrine** | 1.668 | NA | 1.738 | NA | 0.042 | 3 |
| **Respiratory** | 1.429 | NA | 1.291 | NA | 0.011 | 16 |
| **Gastrointestinal** | 1.425 | NA | 1.256 | NA | 0.385 | 3 |
| **Mortality** | 1.314 | NA | 1.293 | NA | 0.089 | 7 |
| **Social Interactions** | 1.208 | NA | 1.259 | NA | 0.172 | 10 |
| **Immunological** | 1.133 | NA | 0.971 | 1.030 | 0.837 | 14 |
| **Ear, Nose, Throat** | 1.103 | NA | 0.904 | 1.106 | 0.829 | 4 |
| **Neurological** | 1.064 | NA | 0.922 | 1.085 | 0.557 | 10 |
| **Muscular** | 1.034 | NA | 1.165 | NA | 0.540 | 3 |
| **Skeletal** | 1.030 | NA | 1.142 | NA | 0.047 | 24 |
| **Nutritional** | 1.014 | NA | 0.770 | 1.299 | 0.212 | 28 |
| **Cardiovascular** | 0.984 | 1.017 | 0.855 | 1.169 | 0.399 | 13 |
| **Metabolic** | 0.951 | 1.052 | 0.813 | 1.230 | 0.179 | 52 |
| **Reproduction** | 0.934 | 1.070 | 0.928 | 1.077 | 0.659 | 12 |
| **Psychiatric** | 0.918 | 1.089 | 0.627 | 1.595 | 0.024 | 67 |
| **Activities** | 0.915 | 1.093 | 0.387 | 2.583 | 0.001 | 66 |
| **Environment** | 0.828 | 1.207 | 0.323 | 3.094 | 0.011 | 31 |
| **Ophthalmological** | 0.707 | 1.415 | 0.360 | 2.778 | 0.030 | 22 |
| **Cognitive** | 0.659 | 1.518 | 0.512 | 1.954 | 0.002 | 11 |

**Table S4.** For each of the phenotypic subchapters, we list the median and mean heritability enrichment. These results are also plotted in Figs. 2B-2E and Fig. S6. For those subchapters which are depleted (enrichment below 1), we also report the fold-depletion (1/Enrichment). Subchapters are ordered by their median enrichment. The mean enrichment was calculated on the log-transformed enrichment values. For enrichments that were calculated to be less than zero by S-LDSC (because of high variability in estimates in regions with low heritability), we assigned them the lowest positive enrichment value (0.000497) before log-transforming to calculate the mean enrichment.

| Subchapter | Enr. (median) | Fold-depletion (median) | Enr. (mean) | Fold-depletion (mean) | P | N (traits) |
|---|---|---|---|---|---|---|
| Functions of Hair | 2.602 | NA | 2.719 | NA | 0.006 | 3 |
| Endocrine Gland Functions | 2.164 | NA | 2.164 | NA | NA | 1 |
| Disorders of Puberty, Not Elsewhere Classified | 2.005 | NA | 1.604 | NA | 0.220 | 3 |
| Superficial Injuries Involving Multiple Body Regions | 1.784 | NA | 1.320 | NA | 0.791 | 2 |
| Control of Voluntary Movement Functions | 1.748 | NA | 1.748 | NA | NA | 1 |
| Vasomotor and Allergic Rhinitis | 1.624 | NA | 1.600 | NA | 0.012 | 3 |
| Structure of Mouth | 1.595 | NA | 1.631 | NA | 0.139 | 4 |
| Diabetes Mellitus | 1.562 | NA | 1.558 | NA | 0.097 | 2 |
| Exercise Tolerance Functions | 1.468 | NA | 1.190 | NA | 0.387 | 7 |
| Menstruation Functions | 1.461 | NA | 1.369 | NA | 0.546 | 2 |
| Diverticular Disease of Intestine | 1.425 | NA | 1.425 | NA | NA | 1 |
| Family Relationships | 1.400 | NA | 1.597 | NA | 0.200 | 3 |
| Asthma | 1.388 | NA | 1.396 | NA | 0.035 | 3 |
| Disorders of Lens | 1.366 | NA | 1.366 | NA | NA | 1 |
| All-Cause Mortality | 1.314 | NA | 1.293 | NA | 0.089 | 7 |
| Structures related to movement, unspecified | 1.311 | NA | 1.232 | NA | 0.004 | 12 |
| Conversation and Use of Communication Devices and Techniques | 1.270 | NA | 1.270 | NA | NA | 1 |
| Unspecified | 1.266 | NA | 1.266 | NA | NA | 1 |
| Problems Related to Upbringing | 1.239 | NA | 1.239 | NA | NA | 1 |
| Mental and Behavioural Disorders Due to Use of Tobacco | 1.235 | NA | 1.208 | NA | 0.093 | 11 |
| Abnormalities of Breathing | 1.223 | NA | 1.223 | NA | NA | 1 |
| WBC | 1.208 | NA | 1.271 | NA | 0.075 | 7 |
| Depressive Episode | 1.194 | NA | 1.232 | NA | 0.250 | 7 |
| Assets | 1.179 | NA | 1.179 | NA | NA | 1 |
| Pain in Chest | 1.161 | NA | 1.135 | NA | 0.661 | 2 |
| Persons with Potential Health Hazards Related to Socioeconomic and Psychosocial Circumstances | 1.147 | NA | 0.912 | 1.096 | 0.799 | 6 |
| Heart Functions | 1.118 | NA | 1.197 | NA | 0.724 | 3 |
| Hearing Functions | 1.103 | NA | 0.904 | 1.106 | 0.829 | 4 |
| Other Arthrosis | 1.053 | NA | 0.848 | 1.179 | 0.849 | 2 |
| Informal Social Relationships | 1.030 | NA | 1.137 | NA | 0.531 | 7 |
| Height | 1.019 | NA | 0.987 | 1.014 | 0.758 | 3 |
| Food | 1.014 | NA | 0.770 | 1.299 | 0.212 | 28 |
| Gonarthrosis [Arthrosis of Knee] | 1.010 | NA | 1.010 | NA | 0.000 | 2 |
| Angina Pectoris | 1.008 | NA | 1.007 | NA | 0.809 | 2 |
| Structure of Ankle | 1.002 | NA | 0.972 | 1.029 | 0.593 | 3 |
| Sensation of Pain | 0.999 | 1.001005 | 0.893 | 1.119 | 0.555 | 7 |
| Weight Maintenance Functions | 0.993 | 1.006834 | 0.974 | 1.026 | 0.390 | 39 |
| Muscle Power Functions | 0.955 | 1.047593 | 0.951 | 1.051 | 0.657 | 2 |
| Blood Pressure Functions | 0.948 | 1.055391 | 0.601 | 1.664 | 0.371 | 4 |
| Acquisition of Necessities | 0.940 | 1.063349 | 0.252 | 3.970 | 0.336 | 6 |
| Looking After One's Health | 0.915 | 1.092941 | 0.395 | 2.530 | 0.004 | 54 |
| Essential (Primary) Hypertension | 0.887 | 1.127904 | 0.887 | 1.128 | NA | 1 |
| Sleep Functions | 0.857 | 1.167432 | 0.805 | 1.242 | 0.394 | 6 |
| Disorders of Gallbladder, Biliary Tract and Pancreas | 0.839 | 1.192415 | 0.839 | 1.192 | NA | 1 |
| Psychomotor Functions | 0.836 | 1.196256 | 0.836 | 1.196 | NA | 1 |
| General Metabolic Functions | 0.816 | 1.226145 | 0.297 | 3.367 | 0.389 | 6 |
| Temperament and Personality Functions | 0.812 | 1.230793 | 0.405 | 2.469 | 0.019 | 35 |
| Chronic Ischaemic Heart Disease | 0.783 | 1.277077 | 0.783 | 1.277 | NA | 1 |
| Education | 0.779 | 1.283499 | 0.097 | 10.276 | 0.098 | 8 |
| Sexual Functions | 0.776 | 1.288244 | 0.770 | 1.298 | 0.284 | 2 |
| Mental and Behavioural Disorders Due to Use of Alcohol | 0.738 | 1.354964 | 0.789 | 1.267 | 0.319 | 8 |

| | | | | | | |
|---|---|---|---|---|---|---|
| **Water, Mineral and Electrolyte Balance Functions** | 0.708 | 1.413285 | 0.702 | 1.425 | 0.216 | 7 |
| **Structure of Eyeball** | 0.688 | 1.453888 | 0.338 | 2.960 | 0.028 | 21 |
| **Sleep Disorders** | 0.685 | 1.459958 | 0.685 | 1.460 | NA | 1 |
| **RBC** | 0.674 | 1.483319 | 0.656 | 1.525 | 0.121 | 6 |
| **Higher-Level Cognitive Functions** | 0.659 | 1.518338 | 0.519 | 1.927 | 0.006 | 9 |
| **Work and Employment** | 0.658 | 1.520355 | 0.528 | 1.894 | 0.124 | 4 |
| **Procreation Functions** | 0.609 | 1.641079 | 0.617 | 1.621 | 0.063 | 5 |
| **Potential Health Hazards Related to Socioeconomic and Psychosocial Circumstances** | 0.581 | 1.720078 | 0.269 | 3.711 | 0.117 | 10 |
| **Moving Around Using Transportation** | 0.512 | 1.952477 | 0.307 | 3.252 | 0.313 | 4 |
| **Memory Functions** | 0.276 | 3.61797 | 0.276 | 3.618 | NA | 1 |

**Table S5.** For 41 traits, we calculated partitioned heritability and direction of effect for the Altai-matching introgressed variants (Set 1, Methods). The first set of columns describes the partitioned heritability results calculated with S-LDSC (enrichment, standard error [SE], and P value). Enrichments above 1 indicate depletion. The second set of columns describes the direction of effect results calculated with SLDP (functional correlation [$r_f$], Z-score, corresponding P value, mu, and mu standard error [see Methods]). Positive Functional correlations and Z-scores indicate a positive relationship with the trait in introgressed variants, whereas negative values indicate a negative relationship with the trait (all with reference to the coding of the GWAS).

| | S-LDSC partitioned $h^2$ | | | SLDP direction of effect | | | | |
|---|---|---|---|---|---|---|---|---|
| **Phenotype** | **$h^2$ Enr** | **SE** | **P** | **$r_f$** | **Z** | **P** | **mu** | **SE(mu)** |
| **AutoimmuneDz** | 3.934 | 1.475 | 0.028 | 6.97E-04 | 1.063 | 0.288 | 2.11E-07 | 2.03E-07 |
| **Balding** | 2.269 | 0.699 | 0.068 | 1.52E-03 | 0.189 | 0.850 | 1.33E-06 | 2E-06 |
| **MenopauseAge** | 2.205 | 1.148 | 0.293 | -9.06E-04 | -1.029 | 0.303 | -5.6E-07 | 5.38E-07 |
| **Sunburn** | 2.078 | 0.865 | 0.208 | 1.82E-03 | 3.171 | 1.5E-3 | 9.94E-07 | 2.98E-07 |
| **HairColor** | 1.935 | 1.277 | 0.465 | -4.79E-04 | -0.627 | 0.531 | -5.3E-07 | 6.59E-07 |
| **SkinColor** | 1.883 | 1.363 | 0.508 | 4.54E-04 | 0.873 | 0.383 | 3.45E-07 | 4.23E-07 |
| **FVC** | 1.819 | 0.448 | 0.069 | -5.37E-04 | -0.155 | 0.877 | -4.4E-07 | 1.08E-06 |
| **Heel_T_Score** | 1.780 | 0.499 | 0.126 | -7.20E-04 | -1.175 | 0.240 | -7.7E-07 | 6.43E-07 |
| **Tanning** | 1.752 | 1.451 | 0.592 | -8.12E-04 | -1.761 | 0.078 | -6.1E-07 | 3.38E-07 |
| **MorningPerson** | 1.625 | 0.458 | 0.174 | 3.28E-04 | 0.591 | 0.554 | 1.8E-07 | 3.02E-07 |
| **Eczema** | 1.544 | 0.742 | 0.459 | -8.00E-04 | -1.447 | 0.148 | -3.8E-07 | 2.61E-07 |
| **EosinophilCount** | 1.528 | 0.414 | 0.202 | -9.39E-04 | -1.249 | 0.212 | -8E-07 | 6.55E-07 |
| **WBCCount** | 1.510 | 0.310 | 0.100 | -1.42E-04 | -0.217 | 0.828 | -1.2E-07 | 5.45E-07 |
| **FEV1_FVC_Ratio** | 1.303 | 0.383 | 0.430 | -6.82E-04 | -1.353 | 0.176 | -6E-07 | 4.36E-07 |
| **DermDz** | 1.204 | 1.390 | 0.884 | -2.30E-05 | -0.033 | 0.974 | -7E-09 | 2.19E-07 |
| **WaistHipRatio** | 1.199 | 0.309 | 0.517 | 6.75E-04 | 0.581 | 0.561 | 4.6E-07 | 6.42E-07 |
| **T2D** | 1.197 | 0.484 | 0.681 | 9.20E-05 | 0.135 | 0.893 | 3.35E-08 | 2.48E-07 |
| **SmokingStatus** | 1.119 | 0.301 | 0.694 | -8.24E-04 | -1.233 | 0.218 | -4.5E-07 | 3.57E-07 |
| **RDW** | 1.114 | 0.394 | 0.772 | 1.09E-03 | 0.766 | 0.444 | 9.21E-07 | 1E-06 |
| **CrohnsDz** | 1.103 | 0.696 | 0.882 | -8.98E-04 | -0.666 | 0.505 | -1.1E-06 | 1.65E-06 |
| **Resp_ENT_Dz** | 1.102 | 0.498 | 0.836 | -5.65E-04 | -1.018 | 0.309 | -2.3E-07 | 2.21E-07 |
| **Height** | 1.028 | 0.272 | 0.918 | -2.00E-04 | -0.371 | 0.710 | -2.9E-07 | 7.65E-07 |
| **NumChildrenBorn** | 0.960 | 0.820 | 0.961 | 1.36E-03 | 0.727 | 0.467 | 4.14E-07 | 5.23E-07 |
| **UC** | 0.933 | 0.760 | 0.929 | -2.96E-04 | -0.292 | 0.771 | -2.9E-07 | 1.03E-06 |
| **YearsOfEd** | 0.915 | 0.236 | 0.718 | 4.35E-04 | 0.551 | 0.581 | 2.78E-07 | 4.99E-07 |
| **BMI** | 0.908 | 0.248 | 0.711 | -6.41E-04 | -1.437 | 0.151 | -5.7E-07 | 4.05E-07 |
| **MenarcheAge** | 0.888 | 0.251 | 0.657 | 7.30E-04 | 0.910 | 0.363 | 6.33E-07 | 6.54E-07 |
| **Schizophrenia** | 0.888 | 0.440 | 0.798 | -2.72E-03 | -3.431 | 6.0E-4 | -3.2E-06 | 8.58E-07 |
| **RBCCount** | 0.878 | 0.310 | 0.692 | 1.48E-03 | 1.364 | 0.172 | 1.33E-06 | 1.13E-06 |
| **SystolicBP** | 0.867 | 0.233 | 0.568 | -9.65E-04 | -1.782 | 0.075 | -7.7E-07 | 4.38E-07 |
| **ASD** | 0.835 | 0.934 | 0.860 | 3.31E-03 | 1.587 | 0.113 | 2.92E-06 | 1.86E-06 |
| **Hypothyroidism** | 0.804 | 0.406 | 0.631 | 1.09E-03 | 1.472 | 0.141 | 4.29E-07 | 2.85E-07 |
| **RA** | 0.683 | 0.906 | 0.725 | 1.35E-03 | 1.422 | 0.155 | 1.04E-06 | 7.07E-07 |
| **PlateletCount** | 0.683 | 0.290 | 0.281 | -5.65E-04 | -1.204 | 0.229 | -6E-07 | 5.57E-07 |
| **Neuroticism** | 0.535 | 0.508 | 0.373 | 1.25E-03 | 0.921 | 0.357 | 7.25E-07 | 7.27E-07 |
| **FirstBirthAge** | 0.477 | 0.470 | 0.270 | -9.79E-04 | -0.913 | 0.361 | -3.8E-07 | 4.02E-07 |
| **LDL** | 0.314 | 0.914 | 0.456 | -6.33E-04 | -0.477 | 0.634 | -2.9E-07 | 6.72E-07 |
| **DepressiveSxs** | -0.027 | 1.042 | 0.336 | -1.08E-03 | -1.016 | 0.310 | -4E-07 | 4.36E-07 |
| **HDL** | -0.177 | 0.706 | 0.102 | 1.18E-03 | 1.038 | 0.299 | 5.58E-07 | 5.65E-07 |
| **HighCholesterol** | -0.322 | 0.457 | 0.006 | -5.09E-04 | -0.911 | 0.362 | -2E-07 | 2.19E-07 |
| **Anorexia** | -0.956 | 1.100 | 0.085 | -9.30E-03 | -4.892 | 1.0E-6 | -5.2E-06 | 1.44E-06 |

**Table S6.** See tab in SupplementalTables.xlsx
For the 8 traits considered by the direction of effect analysis using SLDP (Methods), we identified windows with a strong (Pearson) correlation between Neanderthal LD profile and trait-associated risk or protection (column: window_r). This table contains windows that have at least 15 SLDP regression variants (column: window_numSLDPregressionSNPs), widows that have at least one variant marginally associated with the trait ($P < 1x10^{-4}$, column: window_maxGWASchi2), and windows that overlap at least one Altai-matching Neanderthal introgressed allele (set 1 [Methods]). For each window, we list the overlapping RefSeq protein coding genes. If there are no overlapping genes, we list the 2 closest genes (column: overlapping_or_closest_genes).
